# The binding of OTULIN restrains LUBAC activity to prevent TNF-driven immunopathology

**DOI:** 10.64898/2026.02.27.708452

**Authors:** Wenxin Lyu, Berthe Katrine Fiil, John Rizk, Majken Kjær, Max Benjamin Sauerland, Biao Ma, Malin Frawe Jessen, Rune Busk Damgaard, Mads Gyrd-Hansen

## Abstract

Met1-linked ubiquitin chains (Met1-Ub), synthesised by the linear ubiquitin chain assembly complex (LUBAC) and disassembled by the deubiquitinase OTULIN, critically regulate inflammatory signalling. Although OTULIN’s activity is essential to prevent TNF-driven autoinflammatory pathology and embryonic lethality, the regulatory significance of its direct interaction with LUBAC remains unclear. Here, we reveal that mice harbouring a point mutation (OTULIN^Y56A^) in the OTULIN PUB-interacting motif, which disrupts OTULIN-LUBAC interaction, are viable without spontaneous immunopathology. However, *Otulin*^Y56A/Y56A^ mice exhibited hypersensitivity to TNF-induced toxicity, which was not prevented by inhibiting RIPK1 kinase-mediated cell death. Mechanistically, disruption of the OTULIN-LUBAC interaction led to Met1-linked autoubiquitination, which enhanced LUBAC’s activity and increased Met1-Ub accumulation at the TNF receptor signalling complex. This stabilised the signalling complex even after dissociation from TNF, increased NF-κB signalling and, contrary to loss of OTULIN or its activity, protected cells from TNF-induced apoptosis. During systemic *Listeria monocytogenes* infection, the increased response to TNF in *Otulin*^Y56A/Y56A^ mice exaggerated pathology without affecting bacterial burden. Collectively, we identify the physical association of OTULIN to LUBAC as a critical brake that restricts LUBAC’s function and Met1-Ub-dependent inflammatory signalling, thereby preserving tissue integrity and promoting disease tolerance during acute immune activation.

## Introduction

Inflammation is an essential protective mechanism during infection but can cause tissue damage and fatality when dysregulated^1, 2^. Tumour necrosis factor (TNF) promotes inflammation through activation of mitogen-activated protein kinase (MAPK) and nuclear factor-kappa B (NF-κB) signalling, and indirectly by inducing cell death^2, 3^. The latter exacerbates inflammation by releasing damage-associated molecular patterns (DAMPs) from dying cells^2, 3^. Therefore, appropriate regulation of TNF-driven cytokine production and cell death is critical for preventing sterile inflammation and immune pathology in response to infection.

The linear ubiquitin chain assembly complex (LUBAC), composed of HOIP, HOIL-1 and SHARPIN, is recruited to the TNF receptor 1 (TNFR1) signalling complex (TNF-RSC), where it conjugates Met1-linked ubiquitin (Ub) chains (Met1-Ub) onto various ubiquitinated substrates, including TNFR1^4, 5^. The Met1-Ub functions as a scaffold to facilitate NF-κB signalling and suppresses TNF-induced and RIPK1-mediated cell death through the recruitment or retention of Met1-Ub binding proteins such as the NEMO-IKK complex, A20 and ABIN1/2^5–8^.

Met1-Ub assembly by LUBAC is counterbalanced by the Met1-Ub-specific deubiquitinase (DUB) OTULIN (OTU DUB with linear linkage specificity)^9^. Accordingly, OTULIN dysfunction causes OTULIN-related autoinflammatory syndrome (ORAS), a TNF-driven disease in humans^5, 10–12^. In mice, OTULIN activity is required during embryogenesis by protecting against aberrant cell death and regulating angiogenesis, and OTULIN deficiency or ablation of OTULIN activity in adult mice leads to systemic autoinflammation^10, 13–15^. Mechanistically, OTULIN prevents the accumulation of Met1-Ub on LUBAC subunits and maintains normal LUBAC levels in a cell type-specific manner^10–13^.

LUBAC is also regulated by the DUB CYLD, which preferentially cleaves Lys63-and Met1-Ub^16, 17^. CYLD regulates ubiquitination at receptor signalling complexes and promotes TNF-induced cell death, but its role in regulating Met1-Ub is ambiguous^18, 19^. OTULIN and CYLD (via its adaptor SPATA2) both interact with the HOIP peptide:N-glycanase/UBA- or UBX-containing proteins (PUB) domain via a conserved PUB-interacting motif (PIM) in OTULIN and SPATA2^20, 21^. TNF stimulation rapidly recruits LUBAC in complex with SPATA2-CYLD to the TNF-RSC where CYLD promotes the retention of LUBAC at the TNF-RSC^20^. In contrast, OTULIN appears not to be recruited with LUBAC to receptor signalling complexes^4, 20, 22^. Thus, the physiological role of the OTULIN-LUBAC interaction and how it influences TNF signalling outcomes is not known.^12^

In this study, we reveal that the OTULIN-LUBAC interaction, in contrast to OTULIN activity, is dispensable for embryonic development and immune homeostasis in unchallenged mice. Rather, the interaction restricts LUBAC and limits TNF-induced signalling and cytokine production, which protects against pathology during immune activation.

## Results

### The OTULIN-LUBAC interaction prevents LUBAC auto-ubiquitination but is dispensable for embryonic development

Tyrosine 56 (Y56) within the evolutionarily conserved PIM of OTULIN mediates the binding to the HOIP PUB domain and is essential for OTULIN’s interaction with LUBAC^21, 23^. To investigate the physiological relevance of this interaction, we generated knock-in mice carrying a Y56A point mutation in OTULIN (Figure S1A). Unexpectedly, homozygous *Otulin*^Y56A/Y56A^ mice displayed normal viability (Figure S1B), distinct from the embryonic lethality caused by OTULIN deficiency or catalytic inactivation^10, 13–15^. Genotype distribution of offspring followed Mendelian ratios and adult *Otulin*^Y56A/Y56A^ mice gained body weight (BW) comparably to wildtype (WT) littermates (Figures 1A and 1B). Spleen-to-BW ratios were also unchanged in contrast to mice with myeloid-specific ablation of OTULIN^10^ (Figure S1C). In accordance, flow cytometric analysis showed comparable numbers of myeloid and lymphoid cell populations, as well as similar T cell subset distributions between genotypes (Figures 1C, S1D, and S1E), indicating that *Otulin*^Y56A/Y56A^ mice did not develop spontaneous systemic inflammation as described in adult mice following ablation of *Otulin* or OTULIN activity^10, 13^.

**Figure 1.**
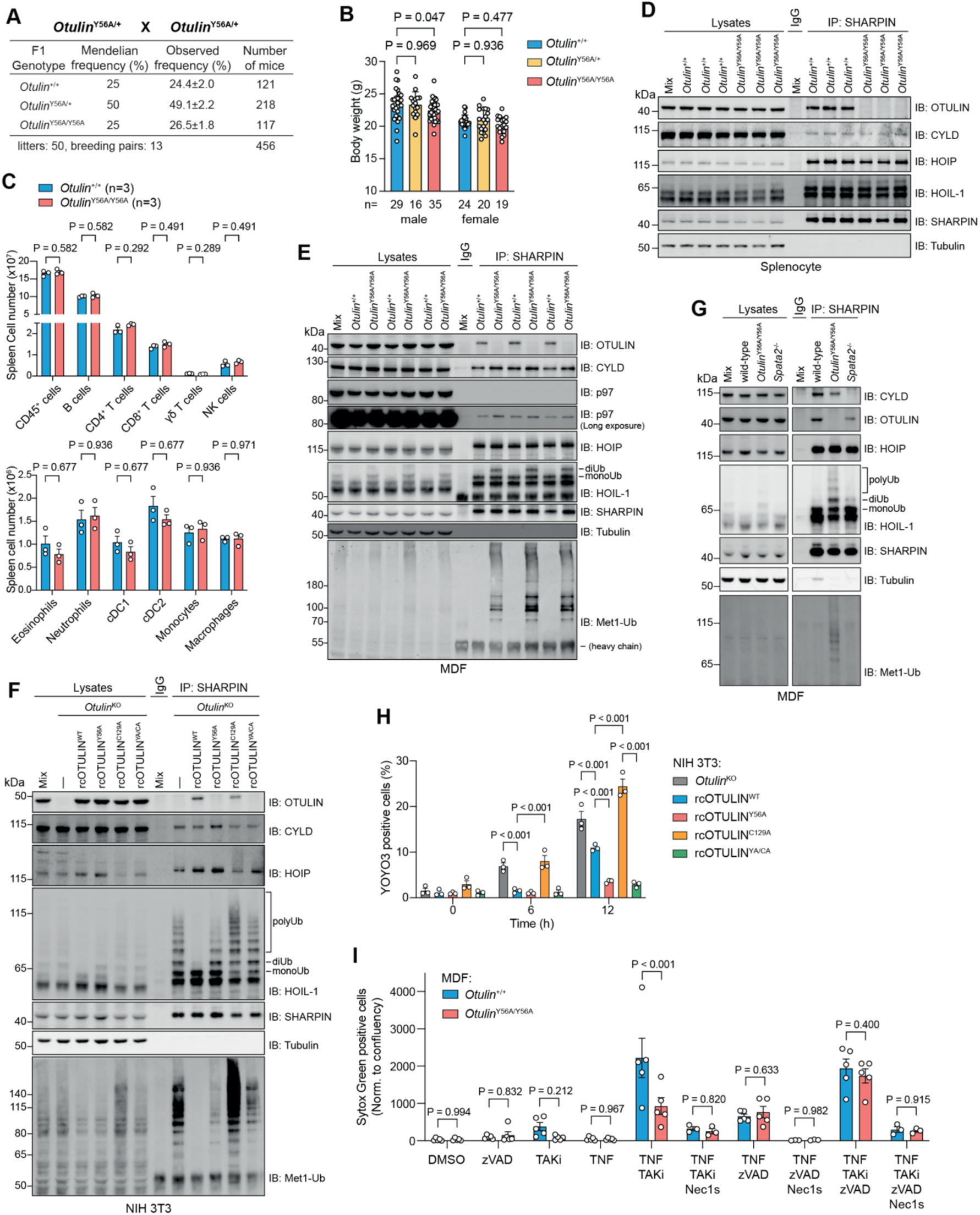
OTULIN-LUBAC interaction is dispensable for embryonic development, immune homeostasis and protection from cell death by TNF despite LUBAC auto-ubiquitination. (A) Observed numbers and ratios of offspring from intercrosses of *Otulin*^Y56A/+^ mice. (B) Body weight of adult male (7-8 weeks old) and female (10-12 weeks old) mice. (C) Total cell number of the indicated immune cell populations in spleen analysed by flow cytometry. (D-G) Immunoblot analysis of immunoprecipitated LUBAC complexes via SHARPIN from splenocytes (D), immortalised MDFs (E, G) or *Otulin*^KO^ NIH 3T3 cells reconstituted with OTULIN variants (F). (H) Cell death of *Otulin*^KO^ NIH 3T3 cells reconstituted with OTULIN variants stimulated with 10 ng/mL mTNF determined by YOYO3 positivity and normalised to cell number. (I) Cell death of primary MDFs pre-treated for 30 minutes with inhibitors (100 nM TAKi, 10 µM Nec1s, 10 µM zVAD or DMSO as vehicle control), followed by stimulation with 10 ng/mL mTNF for 24 hours. Cell death determined by Sytox Green positivity normalised to cell confluence. Data are presented as mean ± SEM, and open circles show data from individual mice or biological replicates. ‘n’ indicates the number of mice or biological replicates in each group. Data in (F, G) are representative of three independent experiments with similar results. Statistical analysis; one-way ANOVA (B), multiple unpaired t-test (C) and two-way ANOVA with Turkey’s multiple comparisons (H, I).

Disruption of the OTULIN-LUBAC interaction in mice was confirmed by immunoprecipitation (IP) of LUBAC from *Otulin*^Y56A/Y56A^ splenocytes, mouse dermal fibroblasts (MDFs) and bone marrow-derived macrophages (BMDMs) (Figures 1D, 1E, and S2A), and by biotin proximity labelling using TurboID coupled to OTULIN that was stably expressed in OTULIN-deficient NIH 3T3 cells^24^ (Figure S2B). TurboID-OTULIN^WT^ extensively biotinylated HOIP, HOIL-1 and SHARPIN but no biotinylation of LUBAC subunits was detected in TurboID-OTULIN^Y56A^ cells (Figure S2B). Notably, the disruption of OTULIN-binding did not affect the interaction of LUBAC with CYLD or with p97, which also contains a PIM that can bind the HOIP PUB domain^21, 23^ (Figures 1D and 1E). Thus, the interaction between OTULIN and LUBAC is dispensable for mouse embryogenesis and immune homeostasis.

Primary *Otulin*^Y56A/Y56A^ cells showed accumulation of Ub-modified HOIL-1 and increased LUBAC-associated Met1-Ub relative to WT cells (Figures 1D, 1E, and S2A). However, this did not lead to a reduction in the level of LUBAC subunits as has been reported in various cell types deficient for OTULIN or OTULIN activity (Figures 1D and 1E)^10–13^. Intrigued by this, we investigated the distinct role of the OTULIN PIM versus loss of OTULIN or OTULIN activity in *Otulin*-knockout NIH 3T3 cells (*Otulin*^KO^) reconstituted with OTULIN variants (Figure 1F). As expected, OTULIN^WT^ efficiently suppressed the elevated Met1-Ub levels and auto-ubiquitination of HOIP and HOIL-1 in OTULIN-deficient cells, and led to increased HOIP and HOIL-1 levels^10,13^ (Figure 1F). OTULIN^Y56A^ also substantially reduced Met1-Ub levels and ubiquitination of HOIP albeit less efficiently than OTULIN^WT^, and increased HOIP and HOIL-1 levels (Figure 1F). However, the Met1-Ub on HOIL-1 was only modestly reduced by OTULIN^Y56A^ relative to OTULIN-deficient cells, in line with the increased ubiquitination of HOIL-1 in *Otulin*^Y56A/Y56A^ MDFs and BMDMs (Figures 1F, 1G, and S2A). In contrast to OTULIN^Y56A^, catalytically inactive OTULIN^C129A^ enhanced Met1-Ub levels and LUBAC auto-ubiquitination and led to a further reduction in HOIP and HOIL-1 relative to the levels in OTULIN-deficient cells, in a manner that was dependent on the interaction with HOIP (Figure 1F). This suggested a) that the OTULIN-HOIP interaction is critical for removing the Met1-Ub conjugated to the auto-monoubiquitinated HOIL-1 (monoUb-HOIL-1) ^25^, and b) that inactive OTULIN bound to LUBAC interferes with the removal of Met1-Ub by other cellular DUBs^21, 26^. We obtained similar results in mouse embryonic fibroblasts (*Otulin*^del/del^ MEFs) reconstituted with the same OTULIN variants although the effects were less pronounced, possibly due to low expression of the reintroduced OTULIN variants (Figure S2C). Treatment of SHARPIN IP samples with recombinant OTULIN confirmed that Ub chains on HOIP and HOIL-1 were Met1-linked (Figure S2C).

CYLD also can disassemble Met1-Ub but disruption of the interaction between LUBAC and CYLD through genetic ablation of *Spata2* did not increase LUBAC auto-ubiquitination or LUBAC-associated Met1-Ub in MDFs (Figure 1G). Also, purification of CYLD-associated LUBAC from OTULIN-mutated MEFs showed clear Met1-Ub-modification of HOIL-1 (Figure S2D). Thus, the OTULIN-HOIP interaction has a non-redundant function in removing Met1-Ub from HOIL-1.

Since ablation of OTULIN or its catalytic activity sensitises to TNF-induced cell death^11, 13, 27, 28^, we next examined if disruption of the OTULIN-LUBAC interaction affected the sensitivity to TNF. OTULIN^Y56A^ protected OTULIN-deficient NIH 3T3 cells and MEFs better or to a similar extent as OTULIN^WT^, whereas OTULIN^C129A^, as reported^13^, accelerated TNF-induced cell death (Figures 1H, S2E, S3A, and S3B). The sensitisation to TNF by OTULIN^C129A^ was reversed by the Y56A mutation (OTULIN^YA/CA^), supporting that disruption of the OTULIN-LUBAC interaction protects from TNF-induced cell death (Figures 1H, S2E, S3A, and S3B). Accordingly, MDFs and BMDMs from *Otulin*^Y56A/Y56A^ mice were less sensitive than WT counterparts to apoptosis when treated with TNF *plus* the TAK1 inhibitor 5Z-7-Oxozeaenol (TAKi) whereas TNF alone did not induce cell death in either genotype (Figures 1I, S3C, and S3D). Necroptotic cell death by TNF in combination with caspase inhibition (zVAD) was induced similarly in both genotypes, which, in line with previous studies^11, 27, 28^, indicates that OTULIN primarily regulates apoptotic cell death (Figures 1I, S3C, and S3D). Inhibition of RIPK1 activity by Necrostatin 2 (Nec1s) largely prevented the cell death of both WT and *Otulin*^Y56A/Y56A^ MDFs (Figures 1I and S3C).

### The OTULIN-LUBAC interaction protects from TNF pathology

OTULIN critically protects from TNF-driven inflammatory pathology^10, 11, 27, 28^. We therefore sought to determine the role of the OTULIN-LUBAC interaction in TNF responses in adult mice. Strikingly, intraperitoneal (i.p.) injection of mouse TNF (mTNF) caused rapid hypothermia within 2-4 hours in *Otulin*^Y56A/Y56A^ mice (Figure 2A), suggestive of systemic inflammatory response syndrome (SIRS)^29^. Hypothermia of *Otulin*^Y56A/Y56A^ mice progressively worsened and they became moribund within hours, whereas *Otulin*^+/+^ and *Otulin*^Y56A/+^ mice remained asymptomatic with minimal reduction in body temperature (Figure 2A).

**Figure 2.**
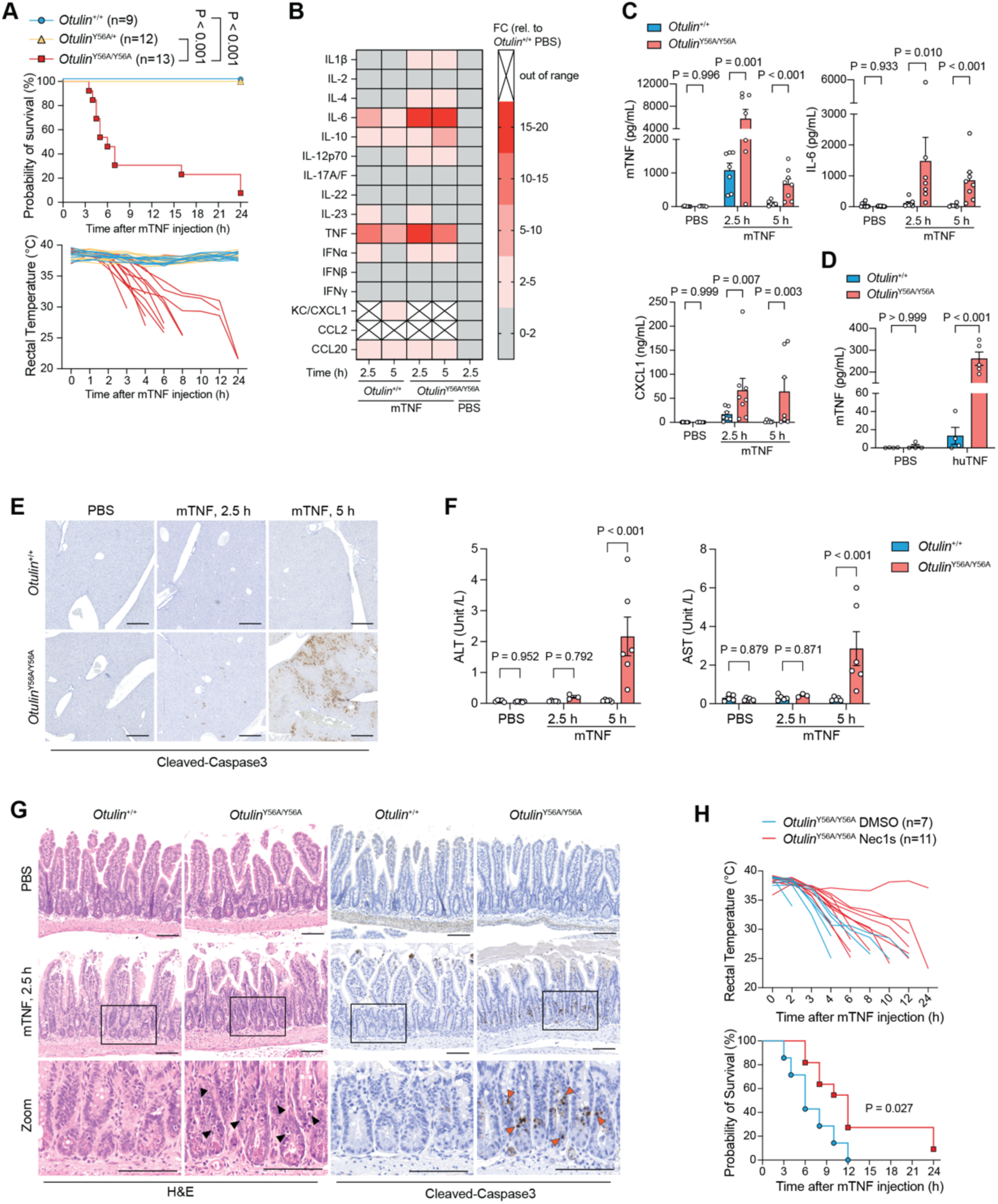
Disruption of the OTULIN-LUBAC interaction sensitises to TNF-induced SIRS. (A) Body temperature and survival of mice injected i.p. with 50 µg/kg mTNF. (B, C, E-G) Mice injected i.p. with PBS or 50 µg/kg mTNF were euthanized after 2.5 h or 5 h and analysed for serum cytokine levels by multiplex cytokine profiling (B) or ELISA (C), H&E or anti-cleaved caspase-3 staining of liver (E) and ileum (G), and serum levels of AST and ALT (F). (D) Serum levels of mTNF measured by ELISA of mice injected i.p. with PBS or 500 µg/kg huTNF euthanized after 2 h. (H) Body temperature and survival of *Otulin*^Y56A/Y56A^ mice injected i.v. with 5% DMSO or 6 mg/kg Nec1s, followed by i.p. injection of 50 µg/kg mTNF. Data are presented as mean ± SEM, and open circles show data from individual mice. ‘n’ indicates the number of mice or biological replicates in each group. Data in (B) represents mean of 3 biological replicates for PBS groups and 5 biological replicates for mTNF groups. Images in (E, G) are representative of three biological replicates with similar results. Arrowheads indicate damaged cells in the crypts. Scale bar: 400 µm (E) and 100 µm (F). Statistical analysis; Log-rank (Mantel-Cox) test (A, H) and two-way ANOVA with multiple comparisons (C, D, F).

Haematological analysis showed a comparable reduction of peripheral white blood cell (WBC) and lymphocyte counts between genotypes, indicating equivalent exposure to mTNF (Figure S4A). *Otulin*^Y56A/Y56A^ mice had a modest but significant increase in red blood cell counts, likely as a result of TNF-driven vascular permeability and fluid loss ^30, 31^ (Figure S4B). In line with this, cytokine profiling revealed elevated levels of pro-inflammatory cytokines, including TNF, IL-6 and CXCL1, in the serum of *Otulin*^Y56A/Y56A^ mice compared with WT littermates prior to the onset of hypothermia (2.5 h; Figures 2B and 2C). To determine if this exacerbated response was mediated by TNFR1 alone or also by TNFR2, mice were administered human TNF (huTNF), which only activates mouse TNFR1^32^. Akin to mTNF, huTNF resulted in heightened cytokine levels in *Otulin*^Y56A/Y56A^ mice relative to WT littermates, which was accompanied by a rapid onset of hypothermia (Figures 2D, S4C, and S4D). Thus, the TNF-hypersensitivity of *Otulin*^Y56A/Y56A^ mice was driven by TNFR1 signalling.

Hepatocyte cell death and liver damage is an early event after i.v. injection of TNF in WT mice^29, 33^. However, only few apoptotic hepatocytes were detected in *Otulin*^Y56A/Y56A^ mice at 2.5 hours after mTNF injection, while widespread cleaved-Caspase-3 positivity was evident by 5 hours (Figure 2E). At this time point livers of *Otulin*^Y56A/Y56A^ mice were visibly darkened, indicative of severe vascular congestion (Figure S4E). Cell death of hepatocytes in *Otulin*^Y56A/Y56A^ mice correlated with significant elevation of serum alanine transaminase (ALT) and aspartate transaminase (AST) levels at 5 hours after mTNF (Figure 2F), indicating that liver pathology occurred after the systemic increase in cytokines and development of hypothermia. WT mice did not show signs of liver damage at either time point (Figures 2E, 2F, and S4E). Paradoxically, *Otulin*^Y56A/Y56A^ mice displayed increased crypt-specific apoptosis of intestinal epithelial cells (IECs) relative to WT littermates already at 2.5 hours after mTNF as evidenced by pyknotic and hyperchromatic nuclei and positive staining for cleaved-Caspase-3^34^ (Figures 2G and S5A). huTNF similarly induced early crypt-specific apoptosis of IECs in *Otulin*^Y56A/Y56A^ mice with little/no liver cell death (Figures S5B and S5C). The localised cell death was reminiscent of mice with ablation of *Otulin* in IECs^28^, albeit less pronounced, which may indicate a distinct role for the OTULIN-LUBAC interaction in protecting crypt IECs in the small intestine against cytotoxic effects of TNF.

RIPK1 activity-mediated cell death is responsible for the development of SIRS in WT mice in response to i.v. injection of TNF ^29, 33^, which prompted us to address if the hypersensitivity of *Otulin*^Y56A/Y56A^ mice to TNF was driven by RIPK1 activity. In accordance with previous reports, Nec1s completely prevented hypothermia and lethality of WT mice in response to i.v. injected TNF^29, 35^ (Figure S5D). However, Nec1s did not prevent lethality of *Otulin*^Y56A/Y56A^ mice following i.p. injection of TNF although it delayed the progression of hypothermia, showing that RIPK1 activity-mediated cell death accelerated the pathology but was not the primary cause of the TNF hypersensitivity (Figure 2H). Together, these data reveal a key role for the OTULIN-LUBAC interaction in protecting against TNF-induced cytokine storm, cell death and tissue damage.

### The OTULIN-LUBAC interaction restricts TNF-induced Met1-Ub and cytokine production

To understand mechanistically how the OTULIN-LUBAC interaction regulates TNF responses, the TNF-RSC was purified using biotin-labelled mTNF (biotin-mTNF). *Otulin*^Y56A/Y56A^ MDFs and BMDMs exhibited substantially more Met1-Ub at the TNF-RSC and more extensive TNFR1 ubiquitination, evident from slow-migrating polyUb-TNFR1 smears, than their WT counterparts (Figures 3A and 3B). This was accompanied by increased abundance of signalling components that rely on Met1-Ub, namely NEMO, IKKβ, TBK1, ABIN-1/2 and A20 (Figures 3A, 3B, S6A, and S6B). The recruitment of LUBAC components and CYLD to the TNF-RSC was also increased in *Otulin*^Y56A/Y56A^ cells relative to WT cells (Figures 3A, 3B, S6A, and S6B). This was despite the elevated auto-ubiquitination of LUBAC in *Otulin*^Y56A/Y56A^ cells, which was further increased in response to TNF, indicating that auto-ubiquitination did not compromise LUBAC’s function in TNF signalling. RIPK1 ubiquitination was comparable between genotypes (Figures 3A, 3B, S6A, and S6B), consistent with the Ub chains on RIPK1 consisting predominantly of linkages other than Met1-Ub^17, 36, 37^. In line with the partial protection of *Otulin*^Y56A/Y56A^ cells from TNF-induced cell death (Figures 1H, 1I, and S3D) and the protective role of the TBK1 and IKK checkpoints^38, 39^, phosphorylation of both kinases was increased at the TNF-RSC and in cell lysates of *Otulin*^Y56A/Y56A^ cells compared with WT cells (Figures 3A, 3B, S6A, and S6B). *Otulin*^Y56A/Y56A^ BMDMs also showed a modest but consistent increase in TNF-induced phosphorylation of IKK-substrates IκBα and the NF-κB subunit RelA relative to WT BMDMs whereas phosphorylation of MAP kinases p38, MK2, ERK1/2 and JNK was comparable between genotypes (Figures 3C and S6C). This translated into an increase in IL-6 and TNF production by IFN-γ or M-CSF primed *Otulin*^Y56A/Y56A^ BMDMs relative to WT BMDMs in response to TNF whereas unprimed BMDMs produced negligible amounts of cytokines (Figures 3D and S6D). The cell surface TNFR1 abundance was comparable between genotypes and was increased substantially by IFN-γ and M-CSF, which likely contributed to the increased cytokine production in both genotypes after priming (Figure S6E).

**Figure 3.**
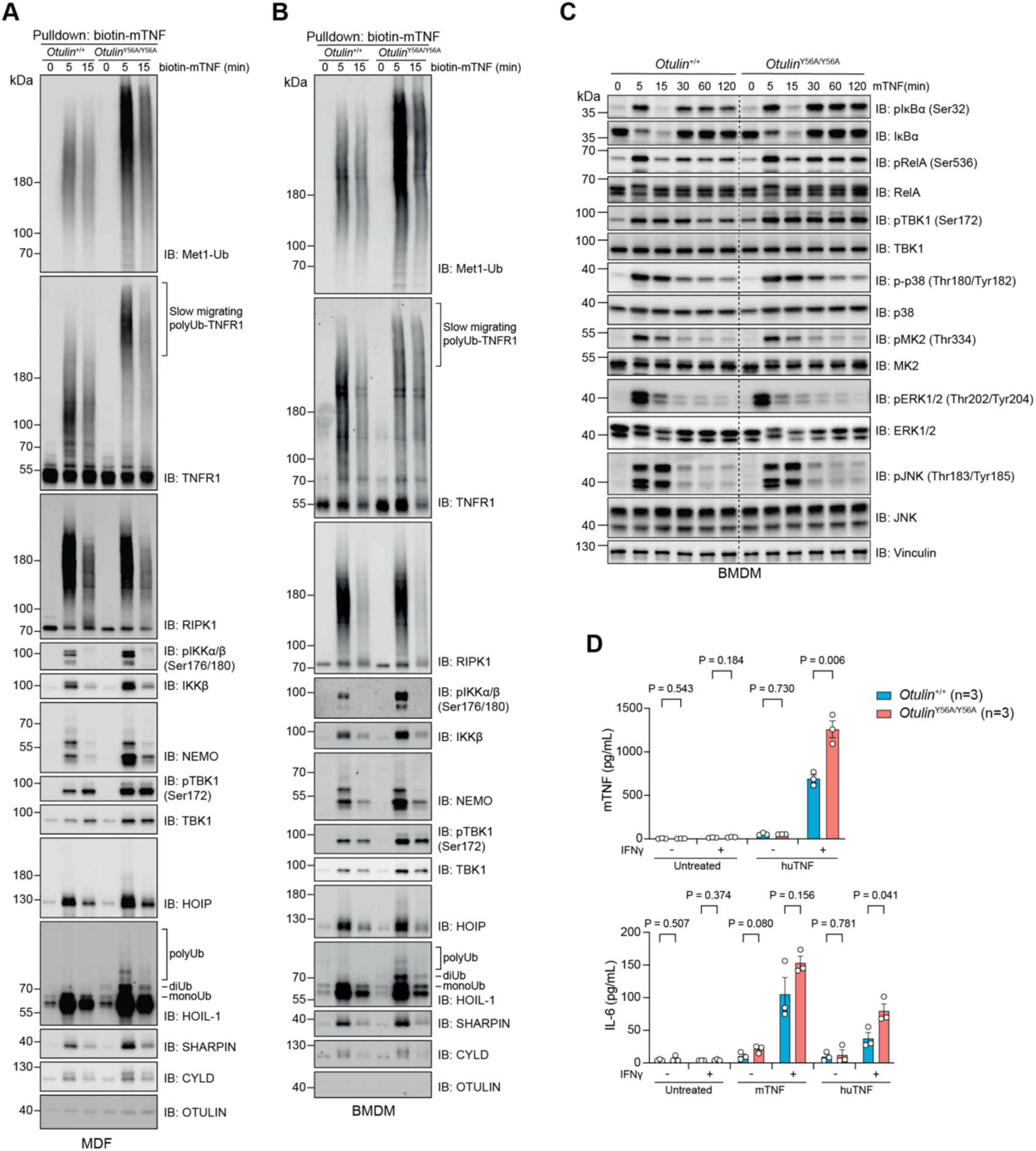
OTULIN-LUBAC interaction restricts TNF-induced Met1-Ub and NF-κB signalling. (A, B) Immunoblot analysis of TNF-RSC purified after stimulation with 50 ng/mL biotin-mTNF for the indicated time points from immortalised MDFs (A) and BMDMs (B). 0 min samples are from cells treated on ice for 2 minutes. (C) Immunoblot analysis of whole cell lysates from BMDMs stimulated with 10 ng/mL mTNF for the indicated time points. (D) Cytokine concentrations in supernatants of BMDMs primed overnight with 20 ng/mL mIFNγ and treated with 10 ng/mL mTNF or 100 ng/mL huTNF for 24 hours. ‘n’ indicates the number of biological replicates in each group. Representative results from three biological replicates (A-C) are shown. Data in (D) are mean ± SEM. Statistical analysis; multiple unpaired *t*-test (D).

Stimulation of IFN-γ-primed BMDMs with the NOD2 ligand L18-MDP showed enhanced TNF and IL-6 production in *Otulin*^Y56A/Y56A^ cells relative WT, in line with the essential role of Met1-Ub in NOD2 signalling^40–42^ (Figure S6F). In contrast, TLR2-and TLR4-mediated cytokine production was comparable between genotypes (Figure S6F). These findings show that the OTULIN-LUBAC interaction restrains the deposition of Met1-Ub at the TNF-RSC (and likely the NOD2 signalling complex) to regulate signalling outcomes by restricting the accumulation of Met1-Ub-dependent signalling factors at the receptor complex.

### Disruption of OTULIN-binding stabilises LUBAC-association with the TNF-RSC

Given the role of Met1-Ub in stabilising the TNF-RSC^43^, we sought to determine if the enhanced Met1-Ub accumulation in *Otulin*^Y56A/Y56A^ cells would affect the disassembly of the TNF-RSC. TNFR1 is internalised within minutes upon TNF sensing, which ultimately leads to disassembly of the complex and is required for the formation of complex II at later timepoints^44–46^. In line with previous studies of the TNF-RSC^19, 43^, we noted a pronounced reduction in signalling complex components within the TNF-bound TNF-RSC at 15 min compared to 5 min after TNF stimulation (Figures 3A and 3B). To track the fate of the signalling complex, we performed sequential purification of the TNF-bound TNF-RSC followed by enrichment of Met1-Ub in the flow-through fraction. This revealed that, in addition to the increased Met1-Ub deposition at the TNF-RSC in *Otulin*^Y56A/Y56A^ cells relative to WT cells, *Otulin*^Y56A/Y56A^ cells also accumulated substantially more Met1-Ub not contained within the TNF-bound TNF-RSC (Figure 4A). This prompted us to analyse the sequential pulldown fractions by mass spectrometry.

**Figure 4.**
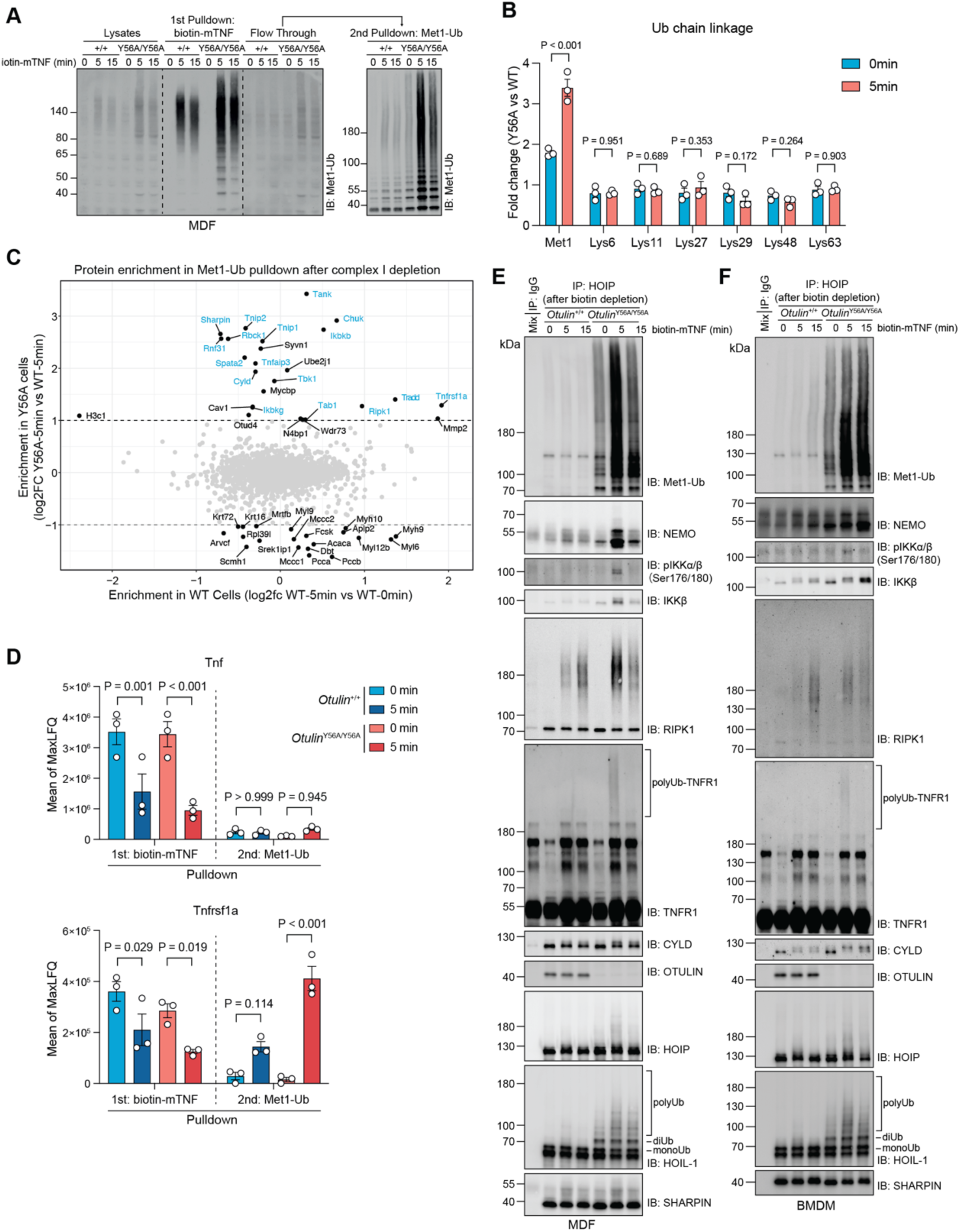
Disruption of OTULIN-LUBAC interaction stabilises the association of LUBAC with TNF-RSC components. (A-D) Immortalised MDFs stimulated with 50 ng/mL biotin-mTNF for the indicated time points. 0 min samples are from cells treated on ice for 2 minutes. Biotin-mTNF pulldowns and Met1-Ub pulldowns from flow-through fractions analysed by immunoblotting (A) or by mass spectrometry-based proteomic analysis of ubiquitin linkages (B), protein enrichment (C), and MaxLFQ intensities (D). (E-F) Immunoblot analysis of immunoprecipitated LUBAC complexes via HOIP from flow-through fraction from samples in (Figures 3A and 3B). Data are presented as mean ± SEM from three biological replicates (B-D). Representative data from three (A, F) or two (E) biological replicates are shown. Statistical analysis; two-way ANOVA with Turkey’s multiple comparisons (B, D).

In accordance with the biochemical data (Figures 3A and 3B), the pulldown of biotin-mTNF enriched most known TNF-RSC components and showed a mild increase of LUBAC subunits and Met1-Ub-associated proteins in samples from *Otulin*^Y56A/Y56A^ cells in comparison with WT cells (Figure S7A). Ubiquitin linkage analysis of the Met1-Ub pulldown from the flow-through fraction revealed a substantial and selective increase of Met1-Ub in *Otulin*^Y56A/Y56A^ cells relative to WT cells, which was further increased by TNF treatment (Figure 4B). This was accompanied by enrichment of most TNF-RSC components, including TNFR1, TRADD, RIPK1, LUBAC, and several proteins recruited via Met1-Ub, namely NEMO-IKK, A20, ABIN-1/2, TBK1, TANK in *Otulin*^Y56A/Y56A^ cells after TNF treatment (Figures 4C, 4D, S7B, and S7C). TNFR1, TRADD and RIPK1 were also enriched in the Met1-Ub pulldown from WT cells but to a lesser degree than in *Otulin*^Y56A/Y56A^ cells (Figures 4C, 4D, and S7C). This increase of TNF-RSC components in *Otulin*^Y56A/Y56A^ cells was not due to residual TNF-bound TNF-RSC since TNF was efficiently captured in the biotin-mTNF pulldown, with no significant difference in TNF abundance in the Met1-Ub pulldown samples (Figure 4D). IP of HOIP following depletion of the TNF-bound TNF-RSC showed that LUBAC associated with several TNF-RSC components as well as Met1-Ub that, in response to TNF, were enriched in *Otulin*^Y56A/Y56A^ cells relative to WT cells (Figures 4E and 4F). Moreover, after TNF treatment HOIP co-purified ubiquitinated forms of TNFR1 and RIPK1 along with NEMO and phosphorylated forms of IKKβ and CYLD, indicating that the complex emanated from the TNF-activated TNF-RSC (Figures 4E, 4F, S7D, and S7E). This showed that the TNF-RSC remains assembled after dissociation from TNF and suggests that disruption of the OTULIN-LUBAC interaction modulates signalling outcomes by stabilising both the TNF-bound and TNF-dissociated TNF-RSC.

### Disruption of OTULIN-binding increases LUBAC activity

Our data posed a conundrum since biochemical purification indicates that OTULIN is not present at the TNF-RSC^20, 22^ (Figures 3A and 3B). In line with this, we did not detect TNF-induced biotinylation of TNFR1 or RIPK1 in cells expressing TurboID-OTULIN (Figure S8A). This implies that the enhanced deposition of Met1-Ub at the TNF-RSC in *Otulin*^Y56A/Y56A^ cells resulted from increased LUBAC function, yet auto-ubiquitination of LUBAC was proposed to be inhibitory^13, 25^.

To directly address how LUBAC auto-ubiquitination in *Otulin*^Y56A/Y56A^ cells influences its enzymatic activity, we performed *in vitro* ubiquitination assays. Endogenous auto-ubiquitinated LUBAC was purified from *Otulin*^Y56A/Y56A^ MDFs by IP of HOIP and was then incubated with USP21 to remove the ubiquitin chains or with buffer as a control (Figure 5A, lane 3 and 4). USP21 treatment led to a clear increase in monoUb-modified HOIL-1, consistent with HOIL-1 depositing the first Ub moiety via an oxyester bond or, alternatively, that the linkage is inaccessible to cleavage by USP21^25, 47^. To rule out that residual USP21 activity in the deubiquitinated samples would interfere Met1-Ub accumulation, USP21-treated LUBAC was incubated with tetra-Met1-Ub, which showed negligible cleavage after 20 min of incubation (Figure 5A, lane 9 and 10). Strikingly, when the ubiquitination reaction was started, the auto-ubiquitinated LUBAC assembled Met1-Ub more efficiently than did USP21-treated LUBAC (Figure 5A). HOIP and HOIL-1 were both extensively ubiquitinated during the reaction but, interestingly, most of HOIL-1 was modified by 10 or less Ub moieties, suggesting that HOIP preferentially extended monoUb or short Ub chains on HOIL-1. This was not due to a general preference for LUBAC to generate short Met1-Ub chains since very slow-migrating Met1-Ub was readily detected (Figure 5A, compare HOIL-1 and Met1-Ub blots). Consistently, inhibition of HOIP activity by Hoipin-8^48^ in OTULIN deficient NIH-3T3 cells reconstituted with OTULIN variants led to a rapid decrease in short Met1-Ub chains on HOIL-1 and in slow-migrating Met1-Ub not conjugated to HOIL-1 (Figures 5B and S8B). Notably, the Hoipin-8 treatment led to a clear increase in monoUb-HOIL-1 in cells with Met1-Ub-modified HOIL-1 and suppressed TNF-induced Met1-Ub accumulation irrespective of the OTULIN status (Figures 5B and S8C). This showed that HOIP activity is required continuously to maintain the Met1-Ub modification of HOIL-1 and that auto-ubiquitination of LUBAC in *Otulin*^Y56A/Y56A^ cells promotes, rather than inhibits, its ability to assemble Met1-Ub. Together, our data suggest that OTULIN, through its interaction with LUBAC, restricts Met1-Ub-mediated TNF signalling by preventing Met1-Ub extension on monoubiquitinated HOIL-1 (and other LUBAC subunits) and thereby limits the propensity of LUBAC to conjugate Met1-Ub on non-LUBAC substrates (Figure 5C).

**Figure 5.**
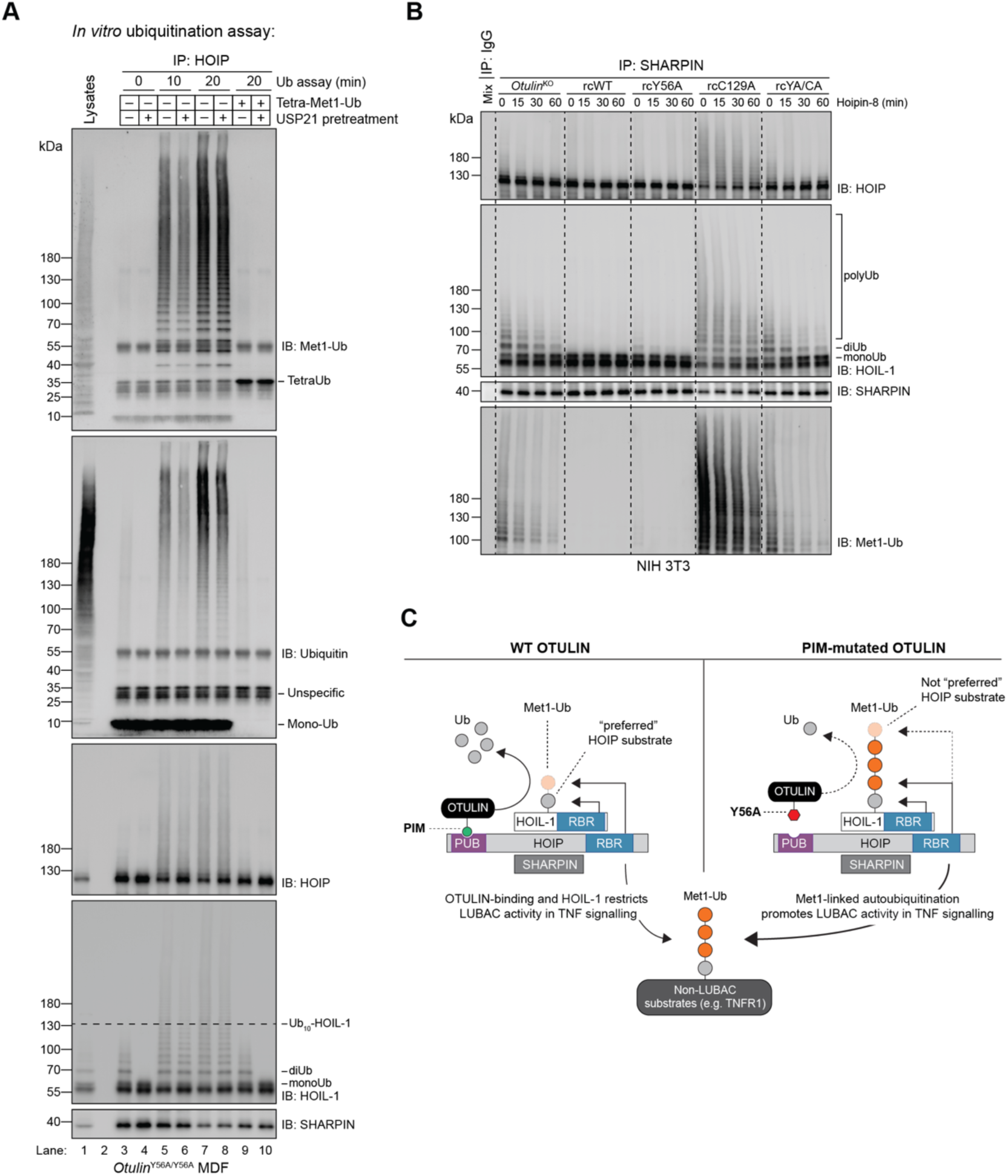
OTULIN counteracts LUBAC auto-ubiquitination to restrict its activity. (A) *In vitro* ubiquitination assays with LUBAC complexes immunoprecipitated via HOIP from immortalised MDFs derived from *Otulin*^Y56A/Y56A^ mice. Samples were incubated with buffer or treated with 1 µM USP21 to remove auto-ubiquitination. Tetra-Met1-Ub was added to samples to determine residual DUB activity. Formation of Met1-Ub was analysed by immunoblotting. (B) Immunoblot analysis of Met1-Ub associated with LUBAC immunoprecipitated via SHARPIN from *Otulin*^KO^ NIH 3T3 cells reconstituted with OTULIN variants treated with 3 µM Hoipin-8 for the indicated time points. (C) Model of how the LUBAC-OTULIN interaction restricts LUBAC activity. Via the PIM-mediated interaction with the HOIP PUB domain, OTULIN counters the Met1-Ub extension on auto-monoubiquitinated HOIL-1 (and possibly other LUBAC subunits) by HOIP in *cis* to maintain HOIL-1 in a mostly auto-monoubiquitinated form, which is a preferred substrate for HOIP. This restricts the propensity of LUBAC to conjugate Met1-Ub on non-LUBAC substrates. Disruption of the OTULIN-LUBAC interaction, e.g. by a Y56A mutation in the OTULIN PIM, enables accumulation of Met1-Ub chains on HOIL-1, limiting the ability of HOIP to further extend the chains. This promotes the generation of Met1-Ub by LUBAC on non-LUBAC substrates in *trans*, leading to enhanced Met1-Ub formation at the TNF receptor signalling complex, increased gene induction, and suppression of apoptosis. Data are representative of three independent experiments with similar results.

### OTULIN-LUBAC interaction protects from systemic pathology in response to *Listeria Monocytogenes* infection

TNF and IFNγ are essential for the early-phase control of *Listeria Monocytogenes* (*Listeria*) infection by macrophages^49–51^. This prompted us to investigate the pathophysiological role of the OTULIN-LUBAC interaction during *Listeria* infection. WT mice challenged with a sublethal dose of *Listeria* experienced a transient drop in BW at day 2 post-infection, which normalised by day 3, whereas *Otulin*^Y56A/Y56A^ mice progressively lost BW (Figure 6A). At this point, haematological analysis indicated impaired host fitness in *Otulin*^Y56A/Y56A^ mice, characterised by a pronounced reduction in WBC, lymphocyte and platelet counts relative to WT mice (Figure 6B). Correspondingly, *Listeria*-infected *Otulin*^Y56A/Y56A^ mice exhibited elevated levels of several proinflammatory cytokines and chemokines relative to WT mice, namely IL-1β, IL-6, IL-17A/F and CXCL1 (Figure 6C). Despite this, bacterial loads in spleen and liver were comparable between WT and *Otulin*^Y56A/Y56A^ mice, suggesting that the ability to control *Listeria* was not compromised by the loss of LUBAC-OTULIN interaction (Figure 6D). Supporting this, *Listeria*-infection of BMDMs *in vitro* showed similar intracellular bacterial growth in WT and *Otulin*^Y56A/Y56A^ BMDMs, which was suppressed equally well by IFN-γ-priming in both genotypes (Figures 6E and S9A)^52,53^. Additionally, infection-induced cell death of BMDMs was similar between genotypes (Figure S9B).

**Figure 6.**
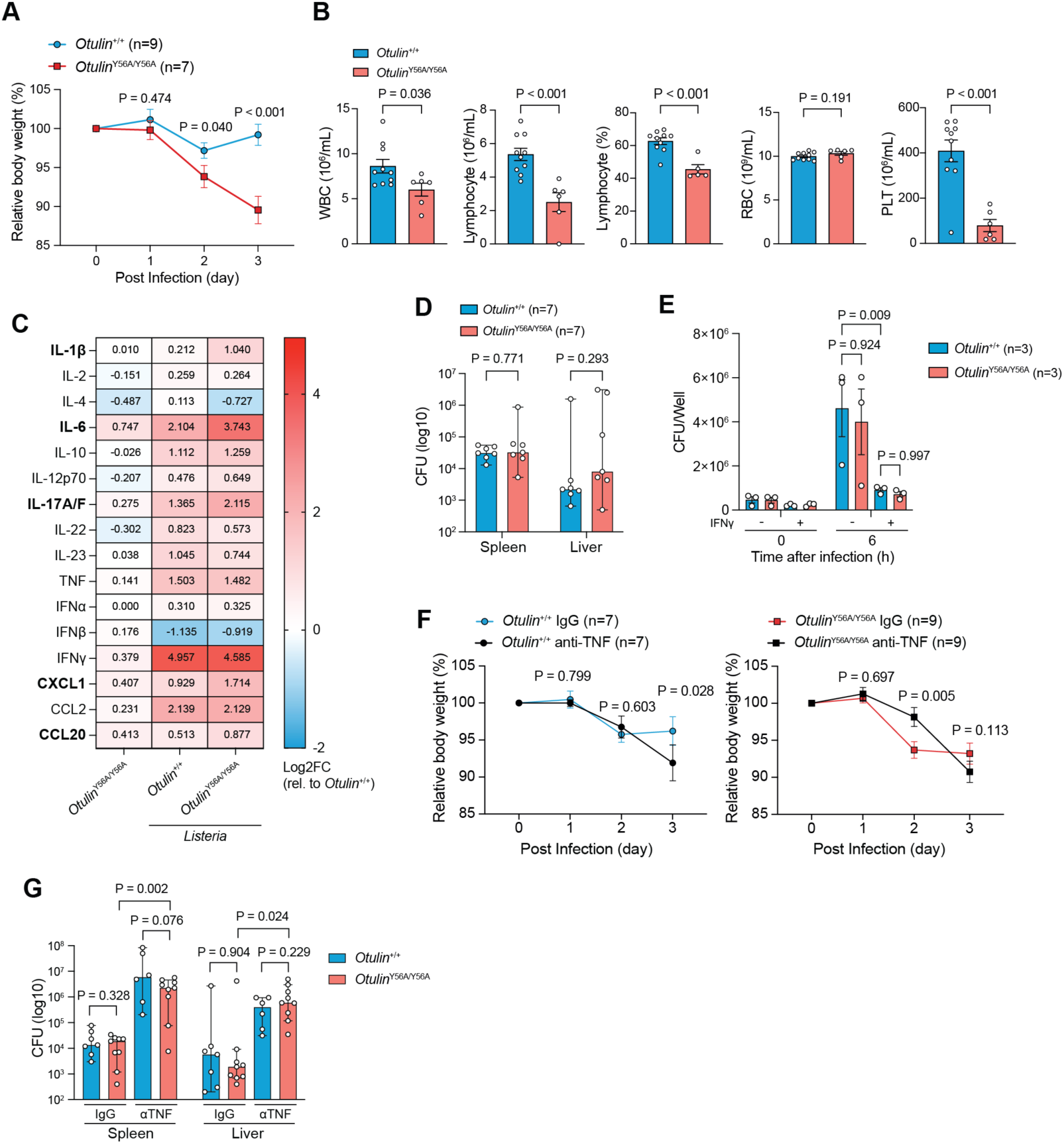
OTULIN-LUBAC interaction protects against *L. monocytogenes* infection-induced pathology driven by TNF. (A) Body weight of mice infected i.p. with 600,000 CFU of *Listeria* monitored daily for 3 days. (B-D) Peripheral blood cell counts (B), Serum cytokine levels (C), and bacterial burden (CFU) in spleen and liver (D) on day 3 of mice in (A). (E) Intracellular bacterial burden (CFU) of BMDMs infected with *Listeria* (MOI=1). BMDMs were primed overnight, or not, with 20 ng/mL mIFN-γ as indicated. (F, G) Body weight (F) and bacterial burden (CFU) in spleen and liver on day 3 (G) of mice infected with *Listeria* as in (A) with or without i.p. injection of 100 µg anti-mTNF antibodies or IgG1 isotype control 12 hours before and 36 hours after infection. Data are presented as mean ± SEM of combined results from two separate experiments, and open circles show data from individual mice. ‘n’ indicates the number of mice in each group. Data in (D, G) are shown as median with 95% confidence intervals. Data in (C) is the mean of 4 biological replicates for Control group and 5 biological replicates for *Listeria* group. Statistical analysis; two-way ANOVA with Turkey’s multiple comparisons (A, E, F) and unpaired two-tailed Student’s *t*-test (B, D, G).

To test if the reduced fitness of *Otulin*^Y56A/Y56A^ mice in response to *Listeria* was caused by TNF-driven pathology, we administered mice with TNF-neutralising antibodies. This significantly improved BW maintenance in *Otulin*^Y56A/Y56A^ mice at day 2 post-infection compared to isotype controls (Figure 6F). However, TNF neutralisation induced abrupt BW loss in both genotypes at day 3 (Figure 6F), which was accompanied by a dramatic accumulation of *Listeria* in spleen and liver, consistent with the critical role of TNF in bacterial control (Figure 6G)^49^. Collectively, this indicates that the OTULIN-LUBAC interaction protects from TNF-driven immunopathology during infection without compromising pathogen control.

## Discussion

The cellular machinery that assembles and disassembles Met1-Ub has emerged as crucially important for regulation of immune responses and inflammation. This is particularly evident for OTULIN whose DUB activity is required to prevent TNF-driven autoinflammatory diseases and lethality of mice during embryogenesis^10, 13–15^.

The fact that *Otulin*^Y56A/Y56A^ mice were viable without signs of spontaneous inflammation indicates that the PIM-PUB interaction between OTULIN and HOIP serves a discrete regulatory function that is mechanistically distinct from OTULIN’s catalytic role in controlling Met1-Ub homeostasis. Supporting this, patients with a biallelic *OTULIN*^R57C^ mutation that attenuates the OTULIN-LUBAC interaction were recently identified with recurrent pyoderma gangrenosum (PG) without the systemic autoinflammatory symptoms characteristic for ORAS patients with mutations that interfere with the DUB activity^54, 55^. Also, ablation of the OTULIN-LUBAC interaction led to a much less pronounced auto-ubiquitination of LUBAC subunits and accumulation of Met1-Ub than observed in cells without OTULIN activity and, importantly, did not result in depletion of LUBAC levels as observed in the absence of OTULIN activity^13^.

RIPK1 kinase activity mediates TNF-induced SIRS in WT mice and the spontaneous TNF-driven inflammatory pathologies described in mice with skin- and liver-specific ablation of *Otulin*^27–29, 56^. In contrast, the exquisite sensitivity of *Otulin*^Y56A/Y56A^ mice to TNF was associated with a rapid increase in serum cytokines levels but was not driven by RIPK1 activity-mediated cell death albeit RIPK1 inhibition delayed the hypothermia. This likely reflects that disruption of the OTULIN-LUBAC interaction promotes NF-kB-driven expression of inflammatory mediators whereas the absence of OTULIN or OTULIN activity sensitises to cell death^11, 13, 27, 28^. Intriguingly, *Otulin*^Y56A/Y56A^ mice displayed early crypt-specific cell death of IECs after TNF challenge which shared features with the intestinal cell death in mice with IEC-specific ablation of *Otulin* or *Tnfaip3* (encodes A20)^28, 34^. Whether this reflects a particular sensitivity of *Otulin*^Y56A/Y56A^ crypt-associated IECs to TNF-induced cell death independently of RIPK1 activity, or an enhanced inflammatory response that disrupted intestinal barrier integrity, will be interesting to delineate in future studies.

*Otulin*^Y56A/Y56A^ mice displayed impaired fitness during *Listeria* infection despite a similar bacterial burden in liver and spleen as their WT counterpart, which points to a critical role for OTULIN-binding in restricting LUBAC-mediated responses to promote disease tolerance during infection by limiting immune-mediated pathology^57^. This appears to be distinct from the role of CYLD during listeriosis as ablation of *Cyld* is shown to reduce the bacterial burden ^58^. Whether this role of CYLD is mediated by its interaction with LUBAC via SPATA2 remains to be determined.

The generation of Met1-Ub by LUBAC in cells with catalytically active OTULIN protects from TNF-induced cell death^2^. This together with the observation that ablation of OTULIN or OTULIN activity leads to extensive Met1-Ub accumulation on LUBAC subunits has led to the notion that auto-ubiquitination inhibits LUBAC function. Our data show that this is not necessarily the case since the auto-ubiquitinated LUBAC in cells with OTULIN^Y56A^ generated more Met1-Ub at the TNF-RSC and protected from cell death similarly or better than LUBAC in cells with WT OTULIN. Moreover, in OTULIN^Y56A^ expressing cells, the association of LUBAC with TNF-RSC components was increased after TNF treatment, which was particularly evident in our analysis of the TNF-bound TNF-RSC followed by enrichment of the LUBAC-associated TNF-RSC. This implicates the OTULIN PIM-mediated interaction with the HOIP PUB domain as a regulatory axis that counterbalances LUBAC activity.

Work by Iwai and colleagues demonstrated that monoubiquitination of LUBAC subunits by HOIL-1 generates a preferred substrate for HOIP to extend Met1-Ub in *cis*, which limits LUBAC-mediated Met1-Ub conjugation on other substrates^25^. This together with our findings suggests that the OTULIN-LUBAC complex is continuously assembling and disassembling auto-Met1-Ub, which suppresses the propensity for LUBAC to assemble Met1-Ub on other substrates. When the OTULIN-LUBAC interaction was disrupted predominantly short HOIP-generated Met1-Ub accumulated on HOIL-1, suggestively because OTULIN^Y56A^ removes the Ub chains less efficiently. An appealing model to explain the increased Met1-Ub activity of auto-ubiquitinated LUBAC is that the Met1-Ub on HOIL-1 become poor substrates for HOIP as the chains are extended, which redirects HOIP to conjugate Met1-Ub in *trans* on substrates such as Ub-modified TNFR1, akin to the scenario without HOIL-1-mediated monoubiquitnation^25^ (Figure 5C). This mechanism may also explain the observation that short Ub chains transiently accumulated on LUBAC subunits in response to TNF as a consequence of recruitment of LUBAC to the TNF-RSC without OTULIN (Figures 3A, 3B, 4E, and 4F)^22^.

In summary, we here uncover an immunoregulatory role for the LUBAC-OTULIN interaction in restricting LUBAC and TNF signalling, which protects tissue integrity during immune activation and promotes host fitness during infection. Further studies of how the interaction is regulated may provide new therapeutic opportunities for inflammatory disorders.

## Supplemental Information

SI document. Materials and Methods, Supplementary Table 1, Supplementary Figures 1-9 and Figure Legends, SI references.

## Acknowledgments

We thank Kit Lee (M.G-H. group) for breeding of mice during the initial phase of the project, members from the M.G-H. and R.B.D. group for helpful advice and suggestions, and Søren Riis Paludan (Aarhus University) for scientific discussion. We acknowledge the valuable contribution to this study made by the Core Facility for Transgenic Animals, the Animal Housing and Breeding Facility at the Department of Experimental Medicine, the Flow Cytometry and Single Cell Core Facility, the Histolab and Veterinary Diagnostic Laboratory at the University of Copenhagen. Mass spectrometry analyses were performed by the Proteomics Research Infrastructure (PRI) at the University of Copenhagen.

## Author Contributions

Conceptualization: MGH, WL

Investigation: WL, BKF

Methodology: WL, BKF, JR, MK, MBS, BM, MFJ

Formal analysis: WL, BKF, MBS

Visualization: WL, BKF, MBS, MGH

Funding acquisition: MGH

Resources: MGH, RBD

Supervision: MG

Writing—original draft: WL, MGH

## Materials & Correspondence

Correspondence relating to the article and material requests should be addressed to Mads Gyrd-Hansen

## Data availability

The mass-spec proteomics data have been deposited to the ProteomeXchange Consortium via the PRIDE^59^ partner repository with the dataset identifier PXD070197.

## REFERENCES

1. Angus Derek, C. & van der Poll, T. Severe Sepsis and Septic Shock. New England Journal of Medicine 369, 840–851.

2. van Loo, G. & Bertrand, M.J.M. Death by TNF: a road to inflammation. Nature Reviews Immunology 23, 289–303 (2023).

3. Huyghe, J., Priem, D. & Bertrand, M.J.M. Cell death checkpoints in the TNF pathway. Trends in Immunology 44, 628–643 (2023).

4. Fiil, B.K. & Gyrd-Hansen, M. The Met1-linked ubiquitin machinery in inflammation and infection. Cell Death & Differentiation 28, 557–569 (2021).

5. Hrdinka, M. & Gyrd-Hansen, M. The Met1-Linked Ubiquitin Machinery: Emerging Themes of (De)regulation. Molecular Cell 68, 265–280 (2017).

6. Ikeda, F. et al. SHARPIN forms a linear ubiquitin ligase complex regulating NF-κB activity and apoptosis. Nature 471, 637–641 (2011).

7. Tokunaga, F. et al. SHARPIN is a component of the NF-κB-activating linear ubiquitin chain assembly complex. Nature 471, 633–636 (2011).

8. Gerlach, B. et al. Linear ubiquitination prevents inflammation and regulates immune signalling. Nature 471, 591–596 (2011).

9. Keusekotten, K. et al. OTULIN antagonizes LUBAC signaling by specifically hydrolyzing Met1-linked polyubiquitin. Cell 153, 1312–1326 (2013).

10. Damgaard, R.B. et al. The deubiquitinase OTULIN is an essential negative regulator of inflammation and autoimmunity. Cell 166, 1215–1230. e1220 (2016).

11. Damgaard, R.B. et al. OTULIN deficiency in ORAS causes cell type-specific LUBAC degradation, dysregulated TNF signalling and cell death. EMBO molecular medicine 11, e9324 (2019).

12. Zhou, Q. et al. Biallelic hypomorphic mutations in a linear deubiquitinase define otulipenia, an early-onset autoinflammatory disease. Proceedings of the National Academy of Sciences 113, 10127–10132 (2016).

13. Heger, K. et al. OTULIN limits cell death and inflammation by deubiquitinating LUBAC. Nature 559, 120–124 (2018).

14. Fu, Y. et al. OTULIN allies with LUBAC to govern angiogenesis by editing ALK1 linear polyubiquitin. Molecular Cell 81, 3187–3204.e3187 (2021).

15. Rivkin, E. et al. The linear ubiquitin-specific deubiquitinase gumby regulates angiogenesis. Nature 498, 318–324 (2013).

16. Komander, D. et al. Molecular discrimination of structurally equivalent Lys 63-linked and linear polyubiquitin chains. EMBO reports 10, 466–473 (2009).

17. Elliott, P.R., et al. Regulation of CYLD activity and specificity by phosphorylation and ubiquitin-binding CAP-Gly domains. Cell Reports 37 (2021).

18. Massoumi, R., Chmielarska, K., Hennecke, K., Pfeifer, A. & Fässler, R. Cyld inhibits tumor cell proliferation by blocking Bcl-3-dependent NF-κB signaling. Cell 125, 665–677 (2006).

19. Wei, R. et al. SPATA2 regulates the activation of RIPK1 by modulating linear ubiquitination. Genes & development 31, 1162–1176 (2017).

20. Elliott, P.R. et al. SPATA2 links CYLD to LUBAC, activates CYLD, and controls LUBAC signaling. Molecular cell 63, 990–1005 (2016).

21. Elliott, P.R. et al. Molecular basis and regulation of OTULIN-LUBAC interaction. Molecular cell 54, 335–348 (2014).

22. Draber, P. et al. LUBAC-recruited CYLD and A20 regulate gene activation and cell death by exerting opposing effects on linear ubiquitin in signaling complexes. Cell reports 13, 2258–2272 (2015).

23. Schaeffer, V. et al. Binding of OTULIN to the PUB Domain of HOIP Controls NF-κB Signaling. Molecular Cell 54, 349–361 (2014).

24. Cho, K.F. et al. Proximity labeling in mammalian cells with TurboID and split-TurboID. Nature Protocols 15, 3971–3999 (2020).

25. Fuseya, Y. et al. The HOIL-1L ligase modulates immune signalling and cell death via monoubiquitination of LUBAC. Nature Cell Biology 22, 663–673 (2020).

26. Davidson, S. et al. Dominant negative OTULIN-related autoinflammatory syndrome. Journal of Experimental Medicine 221 (2024).

27. Hoste, E. et al. OTULIN maintains skin homeostasis by controlling keratinocyte death and stem cell identity. Nature communications 12, 5913 (2021).

28. Verboom, L. et al. OTULIN protects the intestinal epithelium from apoptosis during inflammation and infection. Cell Death & Disease 14, 534 (2023).

29. Duprez, L. et al. RIP Kinase-Dependent Necrosis Drives Lethal Systemic Inflammatory Response Syndrome. Immunity 35, 908–918 (2011).

30. Rivers, E. et al. Early Goal-Directed Therapy in the Treatment of Severe Sepsis and Septic Shock. New England Journal of Medicine 345, 1368–1377.

31. Fajgenbaum, D.C. & June, C.H. Cytokine storm. New England Journal of Medicine 383, 2255–2273 (2020).

32. Lewis, M. et al. Cloning and expression of cDNAs for two distinct murine tumor necrosis factor receptors demonstrate one receptor is species specific. Proceedings of the National Academy of Sciences 88, 2830–2834 (1991).

33. Zelic, M. et al. RIP kinase 1–dependent endothelial necroptosis underlies systemic inflammatory response syndrome. The Journal of Clinical Investigation 128, 2064–2075 (2018).

34. Vereecke, L. et al. Enterocyte-specific A20 deficiency sensitizes to tumor necrosis factor–induced toxicity and experimental colitis. Journal of Experimental Medicine 207, 1513–1523 (2010).

35. Takahashi, N. et al. Necrostatin-1 analogues: critical issues on the specificity, activity and in vivo use in experimental disease models. Cell Death & Disease 3, e437–e437 (2012).

36. Hospenthal, M.K., Mevissen, T.E. & Komander, D. Deubiquitinase-based analysis of ubiquitin chain architecture using Ubiquitin Chain Restriction (UbiCRest). Nature protocols 10, 349–361 (2015).

37. Annibaldi, A. et al. Ubiquitin-mediated regulation of RIPK1 kinase activity independent of IKK and MK2. Molecular cell 69, 566–580. e565 (2018).

38. Lafont, E. et al. TBK1 and IKKε prevent TNF-induced cell death by RIPK1 phosphorylation. Nature Cell Biology 20, 1389–1399 (2018).

39. Dondelinger, Y. et al. NF-κB-independent role of IKKα/IKKβ in preventing RIPK1 kinase-dependent apoptotic and necroptotic cell death during TNF signaling. Molecular cell 60, 63–76 (2015).

40. Damgaard, R.B. et al. The ubiquitin ligase XIAP recruits LUBAC for NOD2 signaling in inflammation and innate immunity. Molecular cell 46, 746–758 (2012).

41. Hrdinka, M. et al. CYLD limits Lys63-and Met1-linked ubiquitin at receptor complexes to regulate innate immune signaling. Cell reports 14, 2846–2858 (2016).

42. Fiil, B.K. et al. OTULIN restricts Met1-linked ubiquitination to control innate immune signaling. Molecular cell 50, 818–830 (2013).

43. Haas, T.L. et al. Recruitment of the linear ubiquitin chain assembly complex stabilizes the TNF-R1 signaling complex and is required for TNF-mediated gene induction. Molecular cell 36, 831–844 (2009).

44. Schütze, S., Tchikov, V. & Schneider-Brachert, W. Regulation of TNFR1 and CD95 signalling by receptor compartmentalization. Nature Reviews Molecular Cell Biology 9, 655–662 (2008).

45. Schneider-Brachert, W. et al. Compartmentalization of TNF receptor 1 signaling: internalized TNF receptosomes as death signaling vesicles. Immunity 21, 415–428 (2004).

46. Schneider-Brachert, W. et al. Inhibition of TNF receptor 1 internalization by adenovirus 14.7 K as a novel immune escape mechanism. The Journal of clinical investigation 116, 2901–2913 (2006).

47. Kelsall, I.R., Zhang, J., Knebel, A., Arthur, J.S.C. & Cohen, P. The E3 ligase HOIL-1 catalyses ester bond formation between ubiquitin and components of the Myddosome in mammalian cells. Proceedings of the National Academy of Sciences 116, 13293–13298 (2019).

48. Katsuya, K. et al. Small-molecule inhibitors of linear ubiquitin chain assembly complex (LUBAC), HOIPINs, suppress NF-κB signaling. Biochemical and Biophysical Research Communications 509, 700–706 (2019).

49. Rothe, J. et al. Mice lacking the tumour necrosis factor receptor 1 are resistant to IMF-mediated toxicity but highly susceptible to infection by Listeria monocytogenes. Nature 364, 798–802 (1993).

50. Unanue, E.R. Inter-relationship among macrophages, natural killer cells and neutrophils in early stages of Listeria resistance. Current opinion in immunology 9, 35–43 (1997).

51. Buchmeier, N.A. & Schreiber, R.D. Requirement of endogenous interferon-gamma production for resolution of Listeria monocytogenes infection. Proceedings of the National Academy of Sciences 82, 7404–7408 (1985).

52. Huang, S. et al. Immune Response in Mice That Lack the Interferon-γ Receptor. Science 259, 1742–1745 (1993).

53. Jessop, F. et al. Interferon Gamma Reprograms Host Mitochondrial Metabolism through Inhibition of Complex II To Control Intracellular Bacterial Replication. Infection and Immunity 88, 10.1128/iai.00744-00719 (2020).

54. Bhattad, S. et al. Profile of 208 patients with inborn errors of immunity at a tertiary care center in South India. Clinical and Experimental Medicine 23, 5399–5412 (2023).

55. Gil, H., Mariskanish, C., Bhattad, S. & Markle, J. Pediatric Pyoderma Gangrenosum in Patients with a Novel Biallelic Mutation in OTULIN. Journal of Human Immunity 1, CIS2025abstract. 2124 (2025).

56. Verboom, L. et al. OTULIN Prevents Liver Inflammation and Hepatocellular Carcinoma by Inhibiting FADD- and RIPK1 Kinase-Mediated Hepatocyte Apoptosis. Cell Reports 30, 2237–2247.e2236 (2020).

57. Medzhitov, R., Schneider, D.S. & Soares, M.P. Disease Tolerance as a Defense Strategy. Science 335, 936–941 (2012).

58. Nishanth, G. et al. CYLD enhances severe listeriosis by impairing IL-6/STAT3-dependent fibrin production. PLoS pathogens 9, e1003455 (2013).

59. Perez-Riverol, Y. et al. The PRIDE database at 20 years: 2025 update. Nucleic acids research 53, D543–D553 (2025).

