## Supplemental Information for "The binding of OTULIN restrains LUBAC activity to prevent TNF-driven immunopathology"

**The supplemental information includes:**

Materials and Methods

Supplementary Table 1

Supplementary Figures 1-9

References

### Materials and Methods

#### Mice

*Otulin*-Y56A conditional knock-in mice (*Otulin*-Y56A$^{flox/flox}$) were generated by Cyagen (www.cyagen.com). Briefly, a cassette containing "loxP-partial intron 1-CDS of exons 2~7-polyA-Rox-Neo cassette-Rox-loxP" was inserted into intron 1, with a point mutation at Tyr56 (TAC to GCC) introduced into the exon 2 within the 3' homology arm. The targeting vector was constructed using bacterial artificial chromosome (BAC) clones, and the Neo cassette was flanked by Rox sites. Gene-targeted ES cells were injected into C57BL/6 blastocysts to generate chimeric mice, which were subsequently intercrossed to obtain homozygous *Otulin*-Y56A$^{flox/flox}$ mice. Correct insertion of the whole cassette was confirmed by Sanger sequencing. Genotyping was performed by PCR using the following primers (Forward: 5'-TTA CTG AAG ACA CCT TGG GAA GTG-3'; Reverse: 5'-ACG GCA ACT AAC TCT GAA CAT GAA-3'). To generate whole-body knock-in mice (*Otulin*$^{Y56A/+}$ mice), *Otulin*-Y56A$^{flox/flox}$ mice were crossed with B6.C-Tg(CMV-cre)1Cgn/J mice. Recombination efficiency was validated using a TaqMan probe assay targeting to the Rox site. Mouse strain Spata2$^{tm1.1(KOMP)Vlcg}$ was obtained from the KOMP Repository and genotyped by PCR using the following primers (Forward 1: 5'-AAG AGC ACA CTA AAT GCA CAC CAC C-3', Reverse 1: 5'-TCT CTC TGT AAT TCC ACC CCT GTG C-3'; Forward 2: 5'-CGT GGC CTG ATT CAT TCC-3', Reverse 2: 5'-ATC CTC TGC ATG GTC AGG TC-3'). All mice used in this study were rederived onto a C57BL/6N background and backcrossed for more than 10 generations. Animals were housed under specific pathogen-free (SPF) conditions at the Department of Experimental Medicine, University of Copenhagen. All animal experiments were conducted in littermates aged between 7-16 weeks, in compliance with licenses granted by the Danish Animal Experiments Inspectorate.

**Primary cells**

L929 cells were cultured in RPMI medium (Gibco, 61870010) supplemented with 10% FBS, 100 U/mL penicillin-streptomycin (Gibco, 15140-122) (hereafter R10 media) for two weeks. The conditioned media was filtered through a 0.45μm filter and stored at –20 °C.

Bone marrow cells were isolated via a centrifugation method as previously described[1]. The collected cells were washed with R10 media and cultured in R10 containing 30 % L929-conditioned media in 15-cm non-coated dishes (Falcon, 351058). After 5-6 days, differentiated macrophages were harvested by trypsinization and seeded for further experiments.

For MDFs, mouse tails were disinfected with iodophor (30 seconds), 70% ethanol (2 minutes), and washed once with MDF media containing DMEM/F12 (Gibco, 31330-095), 15% FBS, 250 ng/mL amphotericin B (Gibco, 15290026), 100 U/mL penicillin-streptomycin, 1 mM sodium pyruvate (Gibco, 11360070) and 50 μM β-mercaptoethanol (Gibco, 31350010). After bone removal, tails were incubated overnight in 4 mg/mL dispase II (Merck, D4693) in MDF media at 4 °C. Afterwards, the skin was washed with MDF media, and dermis was separated from epidermis and cut into small pieces (~1mm). The dermis pieces were placed onto culture dish with fatty tissue facing downward. The skin pieces were covered by 1-2 drops of MDF culture media and incubated at 37 °C for 4-6 hours, followed by adding 9 mL culture media for continuous culture. MDFs migrated out from dermis and became confluent after one week. Cells were passaged once before experimental use. MDFs were immortalized by retrovirus infection carrying SV40T antigen-eGFP-puromycin resistance genes and selected by 1 μg/mL puromycin (Invivo Gen, ant-pr-1).

Splenocytes were isolated by digesting minced spleens in R10 medium containing 0.5 mg/mL collagenase IV (Merck, C5138) and 0.02 mg/mL DNase I (Roche, 10104159001) at 37 °C for 30 minutes with magnetic stirring. The digested tissue was filtered through a 70 μm strainer and treated with RBC lysis buffer (Biolegend, 420301) for 5 minutes on ice. Cells were then washed, filtered through a 40μm strainer, and used for downstream experiments.

**TNF-induced SIRS model**

Recombinant murine TNF (mTNF; Biolegend, 575208) or human TNF (huTNF; Biolegend, 570108) was diluted in PBS and administered to mice via i.p. or i.v. injection at a total volume of 100 μL. Dosages are indicated in the respective figure legends. For necroptosis inhibition, Nec1s (6 mg/kg; Selleck, S8037) was i.v. injected 15 minutes prior to TNF administration.

Following TNF challenge, body temperature was monitored every 1-2 hours using a rectal thermometer with rodent-specific probe (BIOSEB, BIO-TK8851). Mice were euthanized upon reaching humane endpoints (body temperature $< 27$ °C). At designated time points, mice were euthanised by cervical dislocation prior to blood collection and organ harvest.

***Listeria Monocytogenes* infection**

Mice were infected i.p. with 600,000 colony-forming units (CFU) of WT *L. monocytogenes* strain EGD, suspended in 100μL PBS. The remaining bacteria were cultured on brain heart infusion (BHI) agar plates (Thermo, CM1136B) to verify CFU titres every time after injection. Body weight was monitored daily for up to 3 days post-infection. For TNF neutralisation, 100μg anti-mTNF antibody (BioXCell, BP0058) or

IgG1 isotype control (BioXCell, BP0088) diluted in 100µL PBS was administered i.p. 12 hours before and 36 hours after infection. Three days after *L. monocytogenes* infection, mice were euthanised. Liver and spleen were harvested, weighed and placed in ice-cold lysis buffer (0.5% Triton x-100 in PBS). Tissues were homogenised using the TissueRuptor II (Qiagen, 9902755) to release intracellular bacteria. Homogenates were serially diluted in lysis buffer, plated on BHI agar plates at 37°C, and CFU were enumerated after incubation.

For *in vitro* infection, BMDMs were seeded in R10 media without penicillin-streptomycin and incubated overnight to allow full attachment, with or without priming with 20 ng/mL recombinant murine IFN-γ (PeproTech, 315-05). On the following day, WT or GFP strain *L. monocytogenes* were diluted in R10 media without penicillin-streptomycin and added to the macrophages at a multiplicity of infection (MOI) of 1. The culture plates were centrifuged at 1,000 × *g* for 5 minutes and subsequently incubated at 37 °C for 25 minutes. Following infection, the media was replaced with fresh R10 media containing 100 µg/mL gentamicin (Gibco, 15750060) and incubated for 30 minutes to eliminate extracellular bacteria. Cells were either lysed immediately in 0.5% Triton x-100 in PBS (defined as 0-minute control) or further incubated in R10 media containing 20 µg/mL gentamicin for 6 hours prior to lysis. Lysates were diluted serially and plated on BHI agar plates, and CFU were enumerated after incubation at 37 °C overnight.

**Haematological and biochemical analysis**

Peripheral blood cell counts were measured using an automated haematology analyser (Sysmex). For biochemical assays, blood was allowed to coagulate at room temperature, then centrifuged at 2,000 × *g* for 20 minutes. The resulting serum was collected and stored at –80 °C until analysis.

Serum ALT and AST tests were performed at Veterinary Diagnostic Laboratory, University of Copenhagen. Serum levels of TNF, IL-6 and CXCL1 were quantified using ELISA kits following the manufacturers' instructions (Novus Biologicals, DY410, DY406, and DY453). Multiplex cytokine profiling was performed using the MESO QuickPlex SQ 120 (Meso Scale Discovery; MSD), with custom U-plex electrochemiluminescence immunoassays (MSD, K15069M). For samples with undetectable analytes, values were set to the lowest limit of detection.

**Flow cytometry**

Three million splenocytes were washed once by PBS and incubated on ice for 30 minutes with anti-mouse CD16/32 antibody to block Fc receptors. Cells were then washed with FACS buffer (2% FBS in PBS) and stained with antibody cocktails (see Supplementary Table 1) for 30 minutes on ice.

For surface TNFR1 staining, trypsinised cells were washed by PBS, followed by Fc receptor blocking for 30 minutes on ice. Cells were then washed and stained with anti-TNFR1 antibodies or the corresponding isotype control in FACS buffer for 30 minutes on ice.

After staining, cells were washed once, resuspended in 200 µL FACS buffer and filtered through 40 µm filter. 1 µg/mL DAPI (Merck, MBD0015) was added into each sample individually 3 minutes before analysis using a BD LSRFortessa™ X-20 Flow Cytometer. Data analysis was conducted using FlowJo v10 software.

**Cell culture, transfection and retrovirus infection**

NIH 3T3, L929 and Phoenix-ECO cell lines were obtained from the American Type Culture Collection (ATCC). *Otulin*$^{del/flox}$ and *Otulin*$^{del/del}$ MEFs were generated previously[2]. Cells were maintained at 37 °C with 5 % $CO_2$ in appropriate media, following the manufacturer's guidelines, and routinely tested for mycoplasma contamination using the MycoAlert Mycoplasma Detection Kit (Lonza, LZ-LT07-701).

For knocking out OTULIN in NIH 3T3 cells, plasmids co-expressing Cas9, eGFP, and a guide RNA (5'-ACT TCC ATA AGG CGA GTC CG-3') were transfected. GFP-positive cells were single-cell sorted, and monoclonal lines were validated for OTULIN deficiency by immunoblotting, TA cloning (Invitrogen, 450641), and sequencing. OTULIN variants were reconstituted in KO cells by co-transfection of Bac transposon vectors encoding murine OTULIN and piggyBac transposase expression vectors, to generate stable cell lines. Cells were selected with 1 μg/mL puromycin.

In *Otulin*$^{del/del}$ MEFs (puromycin-resistant), reconstitution was performed using Bac transposon vectors co-expressing OTULIN variants and TagBFP2. BFP-positive cells were sorted via BD FACSMelody™ Cell Sorter.

For reconstitution with TurboID-OTULIN, plasmids encoding TurboID fused to the N-terminus of OTULIN were transfected into Phoenix-ECO cells to generate retroviral particles. Viral supernatants were filtered (0.45 μm) and directly used to infect *Otulin*$^{KO}$ NIH 3T3 cells. Positive cells were selected with 1 μg/mL puromycin.

All plasmids were synthesised by VectorBuilder, and transfections were carried out using Lipofectamine™ 3000 Transfection Reagent (Invitrogen, L30000008), according to the manufacturer's protocol.

**Immunoprecipitation**

For immunoprecipitation of endogenous LUBAC complex, splenocytes (50 million cells) or other indicated cell types (~90 % confluence in a 10-cm dish) were washed once with PBS and lysed on ice for 20 minutes in 600 μL co-IP buffer (150 mM NaCl, 25 mM HEPES pH 7.5, 0.5% IGEPAL CA-630, 10% glycerol, 0.5 mg/mL NEM, 1 mM DTT and one tablet each of cOmplete protease inhibitor and PhosSTOP phosphatase inhibitor (Roche) per 10 mL buffer). Whole cell lysates were cleared by centrifugation at 20,000 × *g* for 10 minutes at 4 °C. 50 µL lysate was taken and mixed with 10 µL 6x Laemmli Sample Buffer (LSB; Thermo, J61337-AD) as input controls. The remaining lysate was incubated overnight at 4 °C with 1 µg anti-SHARPIN antibody (ProteinTech, 14626-1-AP) on a rotator, followed by a 3-hour incubation with 10 µL pre-washed Pierce™ Protein G Magnetic Beads (Thermo, 88847). Beads were washed with lysis buffer for 5 times and resuspended in 60 µL 2x LSB. Input and IP samples were denatured at 95 °C for 10 minutes, and 10 µL of each sample was loaded onto 4-12 % Bis-Tris gels (Invitrogen, NuPAGE) and western blot analysis.

**Purification of TNF-induced signalling complexes**

To purify the TNF-bound TNF-RSC, BMDMs (10 million cells in 10-cm dishes) and MDFs (~90 % confluent in 15-cm dishes), were stimulated with 50 ng/mL biotinylated mouse TNF (biotin-mTNF; ACROBiosystems, TNA-M82E9) for the indicated times. For the 0-minute controls, cells were treated with ice-cold PBS followed by a 2-minute incubation with 50 ng/mL biotin-mTNF on ice. Cells were washed once with ice-cold PBS, then lysed in 1 mL TNF lysis buffer (30 mM Tris-HCl pH 7.4, 120 mM NaCl, 2 mM KCl, 2 mM EDTA, 1 % Triton x-100) supplemented with DTT, NEM, protease and phosphatase inhibitor as previously described. Input samples were prepared as above. The remaining lysates were incubated with 15 µL pre-washed Pierce™ Streptavidin Magnetic Beads (Thermo, 88817) overnight at 4 °C on a rotator. Flow-through fractions were collected for subsequent processing, as described in Met1 TUBE pulldown or HOIP IP section. Beads were washed 5 times with lysis buffer and resuspended in 80 µL 2x LSB. Input and pulldown samples were denatured at 95 °C for 10 minutes, and 10 µL of each sample was loaded onto 4-12 % Tris-Glycine gels (Invitrogen, NuPAGE) for western blot analysis.

Flow-through collected after depletion of biotin-mTNF were used at this step to purify Met1-Ub-associated proteins (Met1-Ub pulldown) or LUBAC-associated proteins (HOIP IP). Specifically, 50 µL of flow-through was mixed with 10 µL 6x LSB as input controls. For Met1-Ub pulldown, the remaining samples were incubated overnight at 4 °C with 2 µg biotinylated Met1-Ub TUBE binder (Lifesensors, UM306) and 10 µL pre-washed Streptavidin Magnetic Beads on a rotator. For HOIP IP, the remaining samples were incubated with anti-HOIP antibody (MRC, S174D) for 3 hours, followed by incubation with 10 µL pre-washed Pierce™ Protein G Magnetic Beads for 3 hours. Beads were washed 5 times with lysis buffer and resuspended in 60 µL 2x LSB. Samples were denatured at 95 °C for 10 minutes, and 10 µL of each sample was resolved on 4-12 % Tris-Glycine gels (Invitrogen, NuPAGE) for western blot analysis.

**TurboID proximity labelling assay**

NIH 3T3 cells (~90 % confluent in 6-cm dishes) were treated with 100 µM biotin (Merck, B4501) for the indicated time points. Cells were lysed in 1 mL SDS lysis buffer (0.5% SDS and 0.5% IGEPAL CA-630 in PBS) and centrifuged at 20,000 × *g* for 10 minutes to obtain cleared lysates. 50 µL supernatant was taken and mixed with 10 µL 6x LSB for input controls. The remaining lysate was diluted with lysis buffer to a total volume of 4 mL and desalted using a PD-10 Column (GE Healthcare, GE17-0851-01). The desalted eluate (3.5 mL) was incubated with 25 µL Streptavidin Magnetic Beads overnight at 4 °C on a rotator. Beads were washed with 0.5 % SDS in PBS for three times and biotinylated proteins were eluted in 60 µL 2x LSB at 95 °C for 10 minutes. 20 µL of each sample was loaded onto 4-12 % Bis-Tris gels and analysed by western blot.

**Ubiquitin chain restriction assay**

Following LUBAC immunoprecipitation (described above), beads were washed with lysis buffer for 4 times. Beads were then resuspended in 1 mL PBS and split equally into 2 tubes. The beads were then collected and resuspended in 50 µL DUB buffer (50 mM Tris-HCl pH 7.5, 50 mM NaCl, 1 mM DTT) with or without 1 µM recombinant OTULIN. Samples were incubated at 37 °C while shaking at 600 rpm for 1 hour. Reactions were stopped by placing the tubes on ice and removing the supernatant. Beads were resuspended in 60 µL 2x LSB and denatured at 95 °C for 10 minutes. 10 µL of each sample was loaded onto 4-12 % Bis-Tris gel for western blot analysis.

***In vitro* ubiquitination assay**

Anti-HOIP antibodies (4.5µg) were incubated overnight with 45µL protein G magnetic beads in 500µL co-IP buffer at 4 °C. MDFs derived from *Otulin*$^{Y56A/Y56A}$ mice (~90 % confluence in three 15-cm dishes) were washed once with PBS and lysed on ice for 20 minutes in a total volume of 3mL co-IP buffer supplemented with 1mM DTT and PhosSTOP phosphatase inhibitor. Cell lysates were cleared by centrifugation at 20,000 × *g* for 10 minutes at 4 °C. 30 µL lysate was taken and mixed with 10 µL 6x LSB for input control. The remaining lysate was incubated with the antibody-coated beads for 2 hours at 4 °C on a rotator. Beads were subsequently washed three times with 1 mL co-IP buffer and resuspended in 1 mL DUB buffer. The beads were then split equally into 2 tubes, resuspended in 40 µL DUB buffer with or without 1 µM USP21 (Ubiquigent, 64-0037-050), followed by incubation at 37 °C for 30 minutes to remove USP21-sensitive ubiquitin modifications. After enzymatic digestion, beads were washed 6 times with 1 mL co-IP buffer and twice with Ub reaction buffer (20 mM Tris-HCl pH 7.5, 5 mM $MgCl_2$, 2 mM DTT). Beads were further divided equally into 4 tubes for the ubiquitination assay. Reactions were initiated by adding 30 µL of Ub reaction buffer containing 10 µg/mL UBE1 (Enzo, BML-UW9410), 7.5 µg/mL UbcH7 (Bio-

Techne, E2-640), 100 µg/mL ubiquitin, and 2 mM ATP. Reactions were incubated at 30 °C for the indicated time points. To confirm that residual USP21 does not affect the ubiquitination assay, 30 µL Ub reaction buffer containing 1 µg Met1-linked tetra-ubiquitin (Enzo, BML-UW0785) was added as a control. After incubation, 10 µL 6x LSB was added immediately to terminate the reaction. Samples were denatured at 95 °C for 5 minutes and 10 µL of each sample was loaded onto a 4-12 % Tris-Glycine gel for western blot analysis.

**Immunoblotting**

For preparing whole cell lysates without further immunoprecipitation or pulldown assays, equivalent cells were seeded onto 6-well plates. After stimulation, 250 µL of lysis buffer (1x LSB diluted in 1 % SDS) was added to each well, and collected lysates were boiled at 100 °C for 15 minutes. Equal volumes of the lysates were loaded onto SDS-PAGE gel for separation and subsequently transferred onto nitrocellulose membranes (Cytiva, 10600002). Membranes were blocked with 5% non-fat milk for 1 hour at room temperature, followed by incubation with the indicated primary antibodies, diluted in 1% BSA/PBST, overnight at 4 °C on a shaker. The following day, membranes were incubated with HRP-conjugated secondary antibodies for 1 hour, and protein bands were visualized using a ChemiDoc Imaging System (Bio-Rad) and a chemiluminescence kit (Bio-Rad, 1705062).

Primary and secondary antibodies used for immunoblotting are listed in the Supplementary Table 1.

**Cell death assays**

For cell death assays, 50 µL of culture media was first added to each well of a 48-well plate, followed by seeding 300 µL of cells per well, containing a nucleic acid dye at a final concentration of 250 nM (Sytox Green, Invitrogen, S7020 or YOYO-3, AAT

Bioquest, 156312-20-8). Cells were incubated overnight to allow full attachment. On the following morning, an additional 50 µL of culture media containing the indicated inhibitors was added to cells. After 30-minute incubation, cells were stimulated with 5 µL of cytokines diluted in PBS. Cell images were captured at four different regions of each well every hour using the Incucyte S3 Live-Cell Analysis Instrument (Sartorius). The viability of NIH 3T3 cells was assessed using the Incucyte cell-by-cell module. For MEFs, MDFs, and BMDMs, the number of dead cells per image was normalised to the cell confluency. The following inhibitors were used for cell death assays:

| Drug | Target | Final concen. | Supplier | Cat. Nr. |
|---|---|---|---|---|
| 5Z-7-Oxozeaenol (TAKi) | TAK1 | 100 nM | Merck | O9890 |
| Necrostatin 2 (Nec1s) | RIPK1 | 10 µM | Selleck | S8641 |
| Z-VAD-FMK (zVAD) | Pan-Caspase | 10 µM | Selleck | S7023 |

**Histology and immunohistochemistry**

Tissues fixed in 10 % formalin were embedded in paraffin and sectioned at 4 µm thickness. Haematoxylin and eosin (H&E) staining was performed by the Histolab at University of Copenhagen. For immunohistochemistry (IHC), tissue sections on slides were first baked at 65 °C for 1 hour, then dewaxed in xylene, and rehydrated through a graded series of ethanol. Endogenous peroxidase activity was blocked by incubating the sections in 3 % $H_2O_2$ for 15 minutes. Antigen retrieval was performed by heating the sections at 95 °C to 100 °C for 10 minutes in sodium citrate buffer (pH 6.0) (Akoya, AR6001KT). The sections were then incubated with Blocking One Histo (Nacalai Tesque, 06349-64) at room temperature for 10 minutes. Primary antibodies diluted in background-reducing antibody diluent (Dako, S302281-2) were applied and incubated overnight at 4 °C, followed by incubation with HRP-conjugated secondary antibodies at room temperature for 30 minutes. The slides were developed using the Metal

Enhanced DAB Substrate Kit (Thermo, 34065). The stained slides were imaged using the Axioscan.Z1 (Zeiss) and analysed using QuPath v0.4.3 software.

**Mass spectrometry**

Beads from biotin-mTNF pulldown or Met1-Ub pulldown were washed for 4 times with lysis buffer, followed by 2 times with PBS. During the final wash, the beads were transferred to a new protein low-binding tube.

For LC-MS sample preparation, beads were incubated at 37 °C for 30 minutes with digestion buffer 1 (2 M Urea, 50mM Tris-HCl pH 7.5, 2 mM DTT, 20 μg/mL trypsin). The eluates were then mixed with an equal volume of digestion buffer 2 (2 M Urea, 50 mM Tris-HCl pH 7.5, 10 mM Chloroacetamide) and incubated overnight at room temperature. The next day, tryptic peptide mixtures were acidified to 1% Trifluoroacetic acid (TFA) and loaded onto Evotips (Evosep).

During LC-MS, peptides were separated on a Pepsep 15 cm × 150 μM ID column packed with C18 beads (1.5 μm) using an Evosep ONE HPLC system with the default 30-SPD (30 samples per day) method. The column temperature was maintained at 50 °C. Upon elution, peptides were injected via a CaptiveSpray2 source with 20 μm emitter and analysed on a timsTOF pro2 mass spectrometer (Bruker) operated in PASEF mode. MS data were collected over a 100-1700 m/z range with TIMS mobility range of 0.6-1.6 $1/K_0$. Ion mobility was calibrated using three Agilent ESI-L Tuning Mix ions 622.0289, 922.0097 and 1221.9906. TIMS ramp and accumulation times were set to 100 ms each with 10 PASEF ramps recorded for a total cycle time of 1.17 sec. The MS/MS target intensity and intensity threshold were set to 20,000 and 2,500, respectively. An exclusion list of 0.4 min for precursors within 0.015 m/z and 0.015 V $cm^{-2}$ width was activated.

**Bioinformatic analysis**

Raw LC-MS data were interpreted with FragPipe v22.0, MSFragger v4.1[3-5], IonQuant v1.10.27[6], and Philosopher v5.1[7], using the LFQ-MBR workflow and a canonical mouse fasta file containing common contaminants and 50% decoy sequences. The resulting data were further processed in RStudio v4.3.0 using the msqrob2 workflow[8, 9]. In short, data were log2 transformed, median normalised. Differential expression was calculated via linear modelling with a Benjamini-Hochberg p-value adjustment.

**Quantification and statistical analysis**

Unless otherwise stated, error bars represent the standard error of the mean (SEM). Details of statistical tests, *P* values, and additional parameters are provided in the corresponding figure panels and legends. For pairwise comparisons, variance equality was examined using an F-test and *P* values were calculated using unpaired t-test based on whether the data showed equal or unequal variances. The D'Agostino-Pearson omnibus test was used to assess normality and variance homogeneity. When comparing multiple groups, one-way or two-way ANOVA was applied, with appropriate hoc corrections. Statistical analyses were performed using GraphPad Prism 10. All experiments included a minimum of three biological replicates or were independently repeated at least three times to ensure statistical reliability. Investigators were not blinded to group allocation or data analysis during the study.

### Supplementary Table 1: List of antibodies

#### Immunobloting (WB), flow cytometry (FC), immunohistochemistry (IHC)

| Target & clone | Supplier | Cat. Nr. | Dilution | Application |
|---|---|---|---|---|
| OTULIN | Abcam | ab151117 | 1:1000 | WB |
| OTULIN (E3B8L) | Cell Signaling Technology | 74125 | 1:1000 | WB |
| HOIP | Abcam | ab46322 | 1:1000 | WB |
| HOIP (E6M5B) | Abcam | ab315162 | 1:1000 | WB |
| HOIL-1 (EPR28157-53) | Abcam | ab309104 | 1:1000 | WB |
| SHARPIN | ProteinTech | 14626-1-AP | 1:1000 | WB, IP |
| Met1-Ubiquitin (1E3) | Merck | ZRB2114 | 1:10000 | WB |
| Ubiquitin (P4D1) | Cell Signaling Technology | 3936 | 1:2000 | WB |
| TNFR1 (D3I7K) | Cell Signaling Technology | 13377 | 1:1000 | WB |
| RIPK1 (D94C12) | Cell Signaling Technology | 3493 | 1:1000 | WB |
| Phospho-IKKα/β (Ser176/180) (16A6) | Cell Signaling Technology | 2697 | 1:1000 | WB |
| IKKβ (D30C6) | Cell Signaling Technology | 8943 | 1:1000 | WB |
| NEMO (ERP16629) | Abcam | ab178872 | 1:1000 | WB |
| Phospho-TBK1 (Ser172) (D52C2) | Cell Signaling Technology | 5483 | 1:1000 | WB |
| TBK1 (E8I3G) | Cell Signaling Technology | 38066 | 1:1000 | WB |
| A20 (D13H3) | Cell Signaling Technology | 5630 | 1:1000 | WB |
| ABIN-1 | ProteinTech | 15104-1-AP | 1:1000 | WB |
| ABIN-2 | ProteinTech | 15459-A-AP | 1:1000 | WB |
| Phospho-NF-κB RelA (Ser536) (93H1) | Cell Signaling Technology | 3033 | 1:1000 | WB |
| NF-κB RelA (D14E12) | Cell Signaling Technology | 8242 | 1:1000 | WB |
| Phospho-IκBα (Ser32) (14D4) | Cell Signaling Technology | 2859 | 1:1000 | WB |
| IκBα (44D4) | Cell Signaling Technology | 4812 | 1:1000 | WB |
| IκBα (T.937.7) | Invitrogen | MA5-15132 | 1:1000 | WB |
| CYLD (D1A10) | Cell Signaling Technology | 8462 | 1:1000 | WB, IP |
| CYLD (E10) | Santa Cruz | sc-74435 | 1:1000 | WB |
| p97, VCP (5) | Santa Cruz | sc-57492 | 1:1000 | WB |
| Phospho-SAPK/JNK (Thr183/Tyr185) | Cell Signaling Technology | 9251 | 1:1000 | WB |
| JNK2 (56G8) | Cell Signaling Technology | 9258 | 1:1000 | WB |
| Phospho-ERK1/2 (Thr202/Tyr204) (D13.14.4E) | Cell Signaling Technology | 4370 | 1:1000 | WB |
| ERK1/2 (137F5) | Cell Signaling Technology | 4695 | 1:1000 | WB |
| Phospho-p38 (Thr180/Tyr182) (D3F9) | Cell Signaling Technology | 4511 | 1:1000 | WB |
| p38 | Cell Signaling Technology | 9212 | 1:1000 | WB |
| Phospho-MK2 (Thr334) (27B7) | Cell Signaling Technology | 3007 | 1:1000 | WB |
| MK2 | Cell Signaling Technology | 12155 | 1:1000 | WB |
| Tubulin, hFAB Rhodamine | BioRad | AbD22584 | 1:3000 | WB |
| Vinculin | Sigma | V9131 | 1:3000 | WB |
| β-Actin (13E5) | Cell Signaling Technology | 4970 | 1:3000 | WB |
| Streptavidin, HRP-linked | Abcam | ab7403 | 1:10000 | WB |
| anti-rabbit IgG, HRP-linked | Cell Signaling Technology | 7074 | 1:5000 | WB |
| anti-rabbit IgG, HRP-linked | SouthernBiotech | 4030-05 | 1:10000 | WB |
| anti-mouse igG, HRP-linked | Cell Signaling Technology | 7076 | 1:5000 | WB |

| | | | | |
|---|---|---|---|---|
| anti-mouse igG, HRP-linked | SouthernBiotech | 1030-05 | 1:10000 | WB |
| anti-rabbit IgG light chain, HRP-linked (SB62a) | Abcam | ab99697 | 1:5000 | WB |
| Cleaved Caspase-3 | Cell Signaling Technology | 9661 | 1:400 | IHC |
| Rabbit IgG | Cell Signaling Technology | 2729 | | IP |
| Sheep IgG | R&D | 5-001-A | | IP |
| HOIP (fourth bleed) | MRC PPU reagents | S174D | | IP |

| **Target (clone) & conjugate** | **Supplier** | **Cat. Nr.** | **Dilution** | **Application** |
|---|---|---|---|---|
| B220 (RA3-6B2), AF700 | Invitrogen | 56-0452-82 | 1:100 | FC |
| CD11b (M1/70), APC-eFlour780 | Invitrogen | 47-0112-82 | 1:400 | FC |
| CD11c (N418), PE-Cy7 | Invitrogen | 25-0114-81 | 1:600 | FC |
| CD120a (TNFR Type I) (55R-286), APC | BioLegend | 113006 | 1:100 | FC |
| CD16/32 Fc Block (2.4G2), purified | BD Biosciences | 533142 | 1:100 | FC |
| CD172a SIRPα (P84), BV421 | BD Biosciences | 740071 | 1:200 | FC |
| CD19 (1D3), AF700 | BD Biosciences | 557958 | 1:100 | FC |
| CD3e (145-2C11), PE-CF594 | BD Biosciences | 562286 | 1:100 | FC |
| CD4 (GK1.5), FITC | BD Biosciences | 553729 | 1:200 | FC |
| CD44 (IM7), V500 | BD Biosciences | 560781 | 1:200 | FC |
| CD45 (30-F11), BV480 | BD Biosciences | 566095 | 1:200 | FC |
| CD45 (30-F11), BUV395 | BD Biosciences | 564279 | 1:200 | FC |
| CD62L (MEL-14), APC | BD Biosciences | 561919 | 1:200 | FC |
| CD64 (X54-5/7.1), PE | Invitrogen | 12-0641-82 | 1:200 | FC |
| CD8a (53-6.7), BUV737 | BD Biosciences | 612759 | 1:200 | FC |
| F4/80 (T45-2342), BUV395 | BD Biosciences | 565614 | 1:200 | FC |
| Armenian Hamster IgG Isotype (HTK888), APC | BioLegend | 400912 | 1:100 | FC |
| I-A/I-E (M5/114.15.2), Alexa Fluor 488 | BD Biosciences | 562352 | 1:200 | FC |
| Ly6C (AL-21), APC | BD Biosciences | 560595 | 1:200 | FC |
| Ly6G (IA8), BUV737 | BD Biosciences | 741813 | 1:400 | FC |
| NK1.1 (PK136), AF700 | BD Biosciences | 560515 | 1:100 | FC |
| NK1.1 (PK136), APC-Cy7 | BD Biosciences | 560618 | 1:100 | FC |
| TCR β (H57-597), AF700 | Invitrogen | 56-5961-82 | 1:100 | FC |
| TCR β (H57-597), PE-Cy7 | BD Biosciences | 560729 | 1:200 | FC |
| TCR γδ (GL3), BV421 | BD Biosciences | 562892 | 1:200 | FC |
| SiglecF (E50-2440), BV711 | BD Biosciences | 740764 | 1:200 | FC |
| TER-119 (TER-119), AF700 | Invitrogen | 56-5921-82 | 1:100 | FC |
| XCR1 (ZET), BV650 | BioLegend | 148220 | 1:200 | FC |

### FIGURES and LEGENDS

#### Suppl. Figure S1

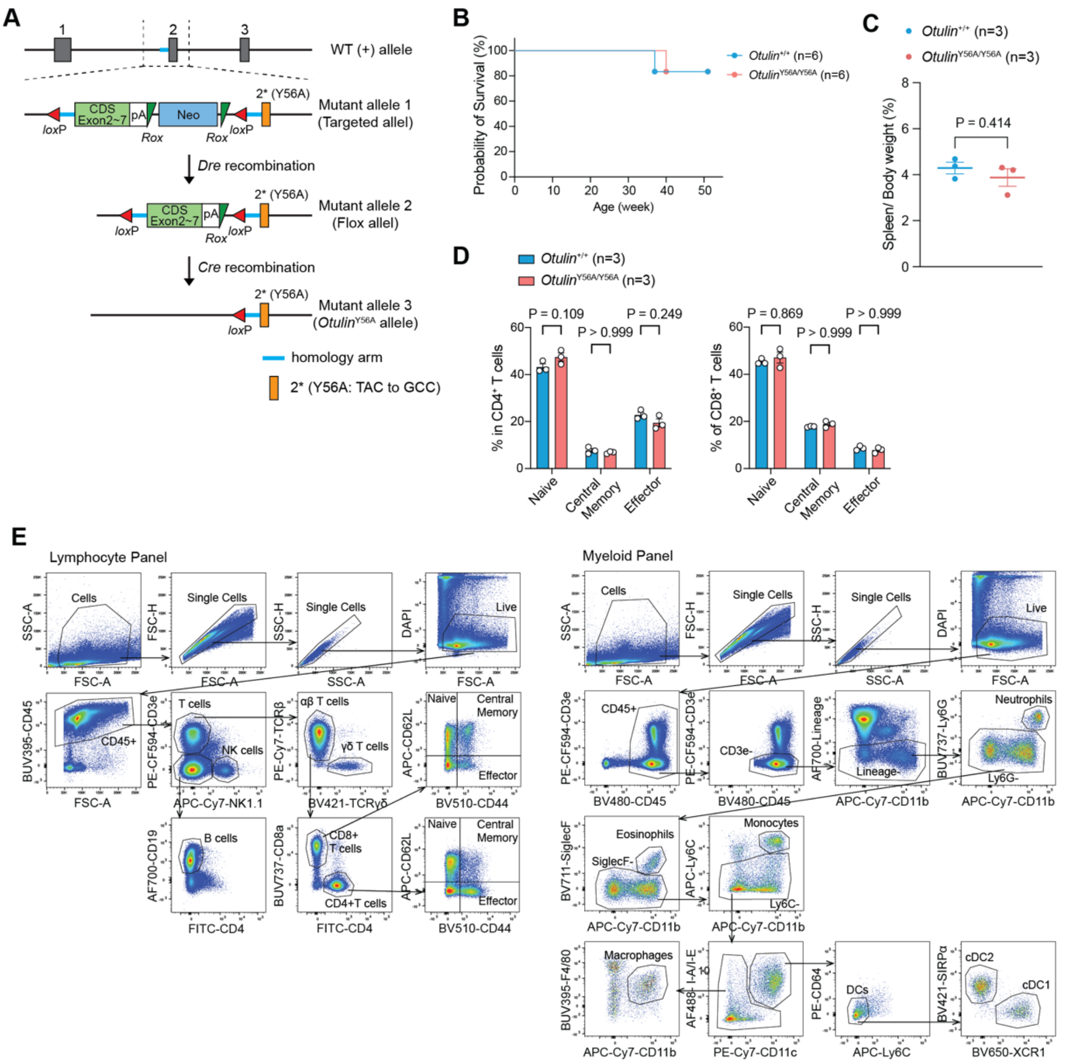


**Suppl. Figure S1. Generation and characterisation of *Otulin*$^{Y56A/Y56A}$ mice.**
(A) Schematic illustration of targeting strategy for *Otulin*$^{Y56A/Y56A}$ mouse generation.
(B, C) Kaplan-Meier plot of survival (B) and spleen-to-body weight ratios (C) for mice of the indicated genotypes.
(D) Proportions of $CD4^+$ and $CD8^+$ T cell subpopulations analysed by flow cytometry.
(E) Gating strategy used in spleen flow cytometry. 'AF700-Lineage' includes AF700-conjugated anti-mouse NK1.1, B220, CD19, TCRβ, and Ter-119 antibodies. Data are presented as mean ± SEM from one experiment with three biological replicates (C, D). Data from individual mice is indicated by circles in graphs. 'n' indicates the number of mice in each group. Statistical analysis; unpaired *t*-test (C) and two-way ANOVA with Bonferroni's multiple comparisons (D).

**Suppl. Figure S2**

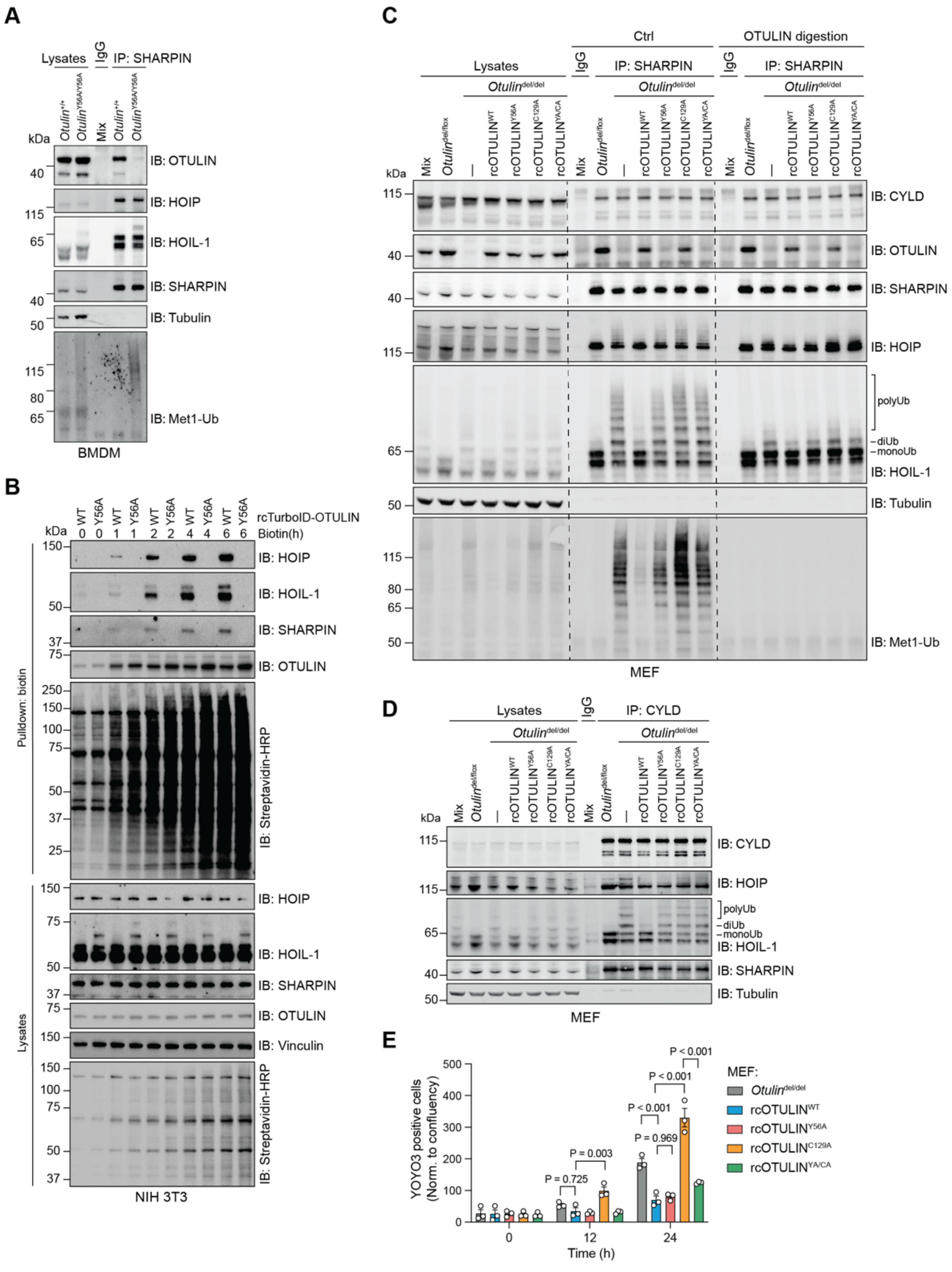


**Suppl. Figure S2. OTULIN$^{Y56A}$ mutation disrupts the OTULIN-LUBAC interaction and leads to LUBAC auto-ubiquitination.**

(A) Immunoblot analysis of LUBAC complexes immunoprecipitated from lysates of BMDMs of indicated genotypes.

(B) Immunoblot analysis of biotinylation of proteins by TurboID-fused OTULIN variants stably expressed in *Otulin*$^{KO}$ NIH 3T3 cells. 100 µM biotin was added for the indicated time points prior to pulldown of biotinylated proteins.

(C) Immunoblot analysis of lysates and LUBAC complexes immunoprecipitated via SHARPIN from immortalised *Otulin*$^{del/flox}$ and *Otulin*$^{del/del}$ MEFs reconstituted with the indicated OTULIN variants. IP samples were incubated, or not, with OTULIN to remove Met1-Ub.

(D) Immunoblot analysis of immunoprecipitated CYLD and co-purified proteins from MEFs.

(E) Cell death of *Otulin*$^{del/del}$ MEFs reconstituted with OTULIN variants after treatment with 10 ng/mL mTNF measured by YOYO3 positivity and normalised to cell confluence. Data shown are one representative experiment of five biological replicates (A) or three independent experiments (B-D) with similar results, and combined results from three independent experiments (E). Data are presented as mean ± SEM and individual values are indicated by open circles (E). Statistical analysis; two-way ANOVA with Turkey's multiple comparisons (E).

**Suppl. Figure S3**

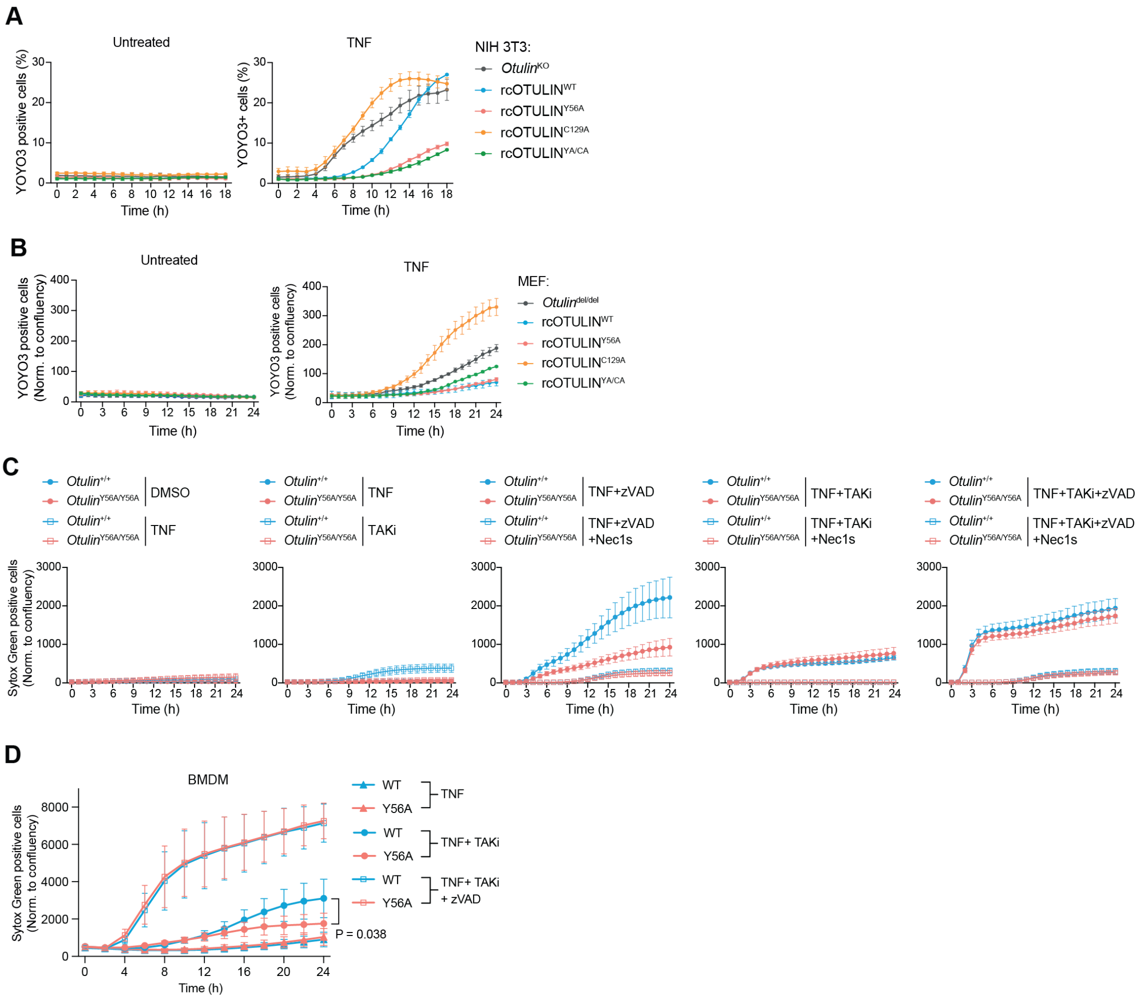


**Suppl. Figure S3. OTULIN$^{Y56A}$ mutant cells are more resistant to TNF-induced apoptosis.**

(A ,B) Cell death of *Otulin*$^{KO}$ NIH 3T3 cells (A) and *Otulin*$^{del/del}$ MEFs (B) reconstituted with OTULIN variants after treatment with 10 ng/mL mTNF measured by YOYO3 positivity and normalised to total cell counts (A) or cell confluence (B).

(C, D) Cell death of primary MDFs (C) and BMDMs (D) treated for 30 minutes with inhibitors as indicated (100 nM TAKi, 10 µM Nec1s, 10 µM zVAD or DMSO as vehicle control) and stimulated with 10 ng/mL mTNF. Cell death determined by Sytox Green positivity and normalised to cell confluence.

Data are presented as mean ± SEM from three independent experiments (A, B), or five (C) or four (D) biological replicates.

**Suppl. Figure S4**

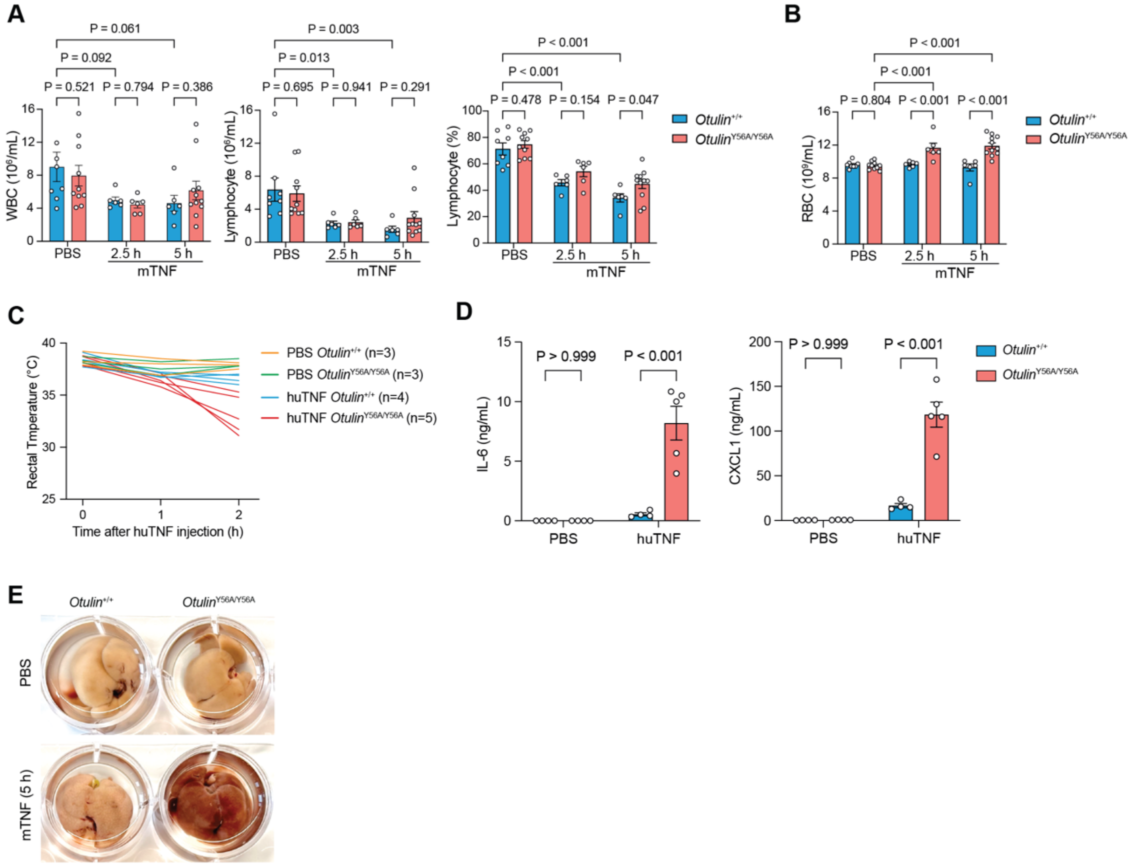


**Suppl. Figure S4. *Otulin*$^{Y56A/Y56A}$ mice are sensitised to TNF-induced SIRS.**
(A, B) Peripheral blood cell counts from mice injected i.p. with PBS for 2.5 hours or with 50 µg/kg mTNF for 2.5 or 5 hours.
(C) Body temperature of mice i.p. injected with 500 µg/kg huTNF.
(D) Serum levels of IL-6 and CXCL1 from mice 2 hours after i.p. injection with 500 µg/kg huTNF.
(E) Livers collected 5 hours post-injection with 50 µg/kg mTNF. Data are presented as mean ± SEM, and open circles show data from individual mice. ‘n’ indicates the number of mice or biological replicates in each group.
Data in (E) shows representative images from three biological replicates. Statistical analysis; two-way ANOVA with Turkey’s multiple comparisons (A, B, D).

**Suppl. Figure S5**

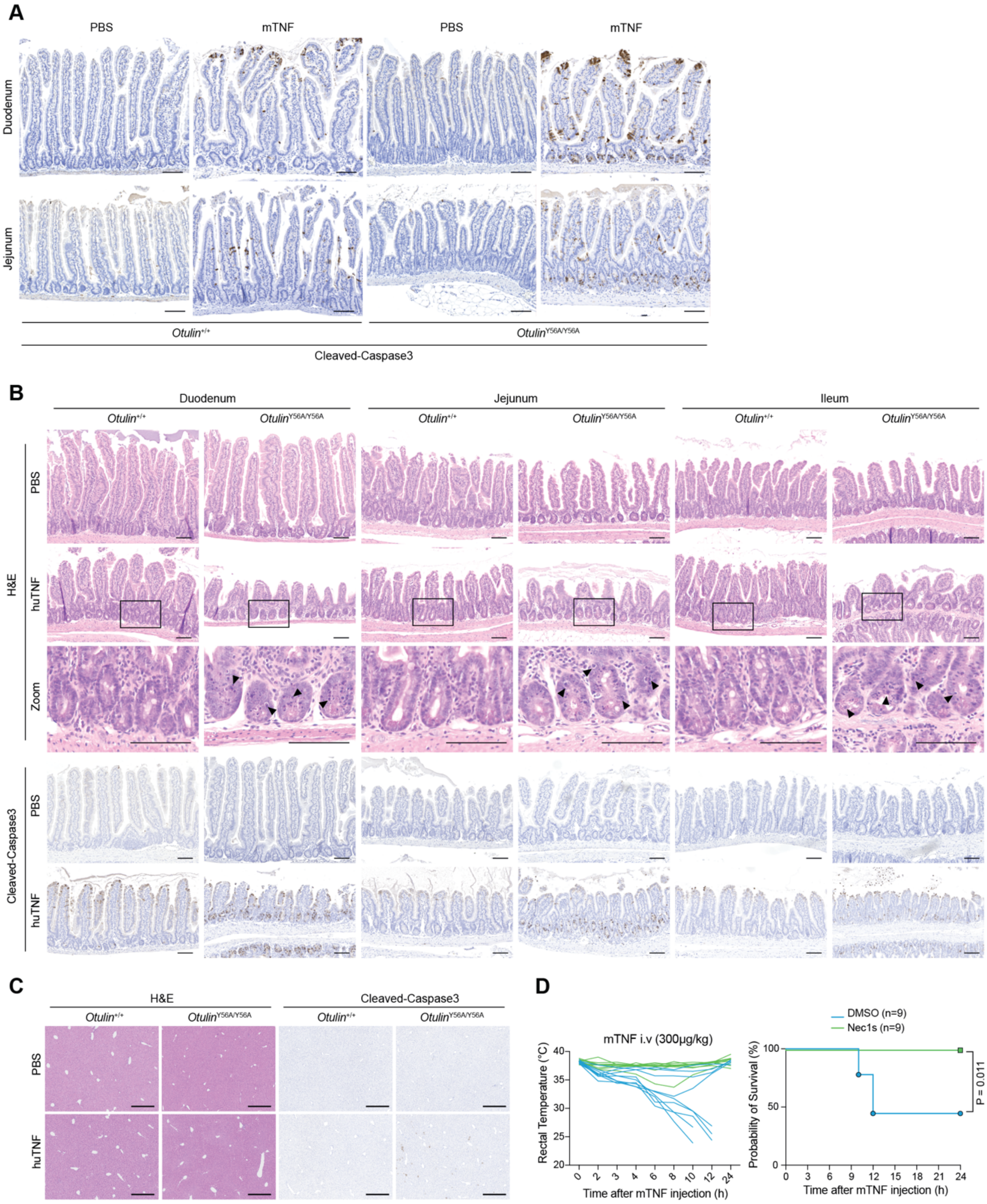


**Suppl. Figure S5. TNF hypersensitivity in *Otulin*$^{Y56A/Y56A}$ mice depends on TNFR1 signalling.**

(A-C) H&E or anti-cleaved caspase-3 staining of the small intestines (A, B) and livers (C) from mice of the indicated genotypes injected i.p. with PBS or 50 µg/kg mTNF and euthanised after 2.5 hours (A) or i.p. injected with PBS or 500 µg/kg huTNF and euthanised after 2 hours (B, C). Arrowheads indicate damaged cells in the crypts (B). Scale bar: 100 µm (A, B), 400 µm (C).

(D) Body temperature and survival of WT mice i.v. injected with 5% DMSO or 6mg/kg Nec1s, followed by i.v. injection of 300 μg/kg mTNF. Body temperature was monitored every 1-2 hours. ‘n’ indicates the number of mice used in each group. Data are presented as one representative image from three biological replicates (A-C). Statistical analysis; Log-rank (Mantel-Cox) test (D).

Suppl. Figure S6

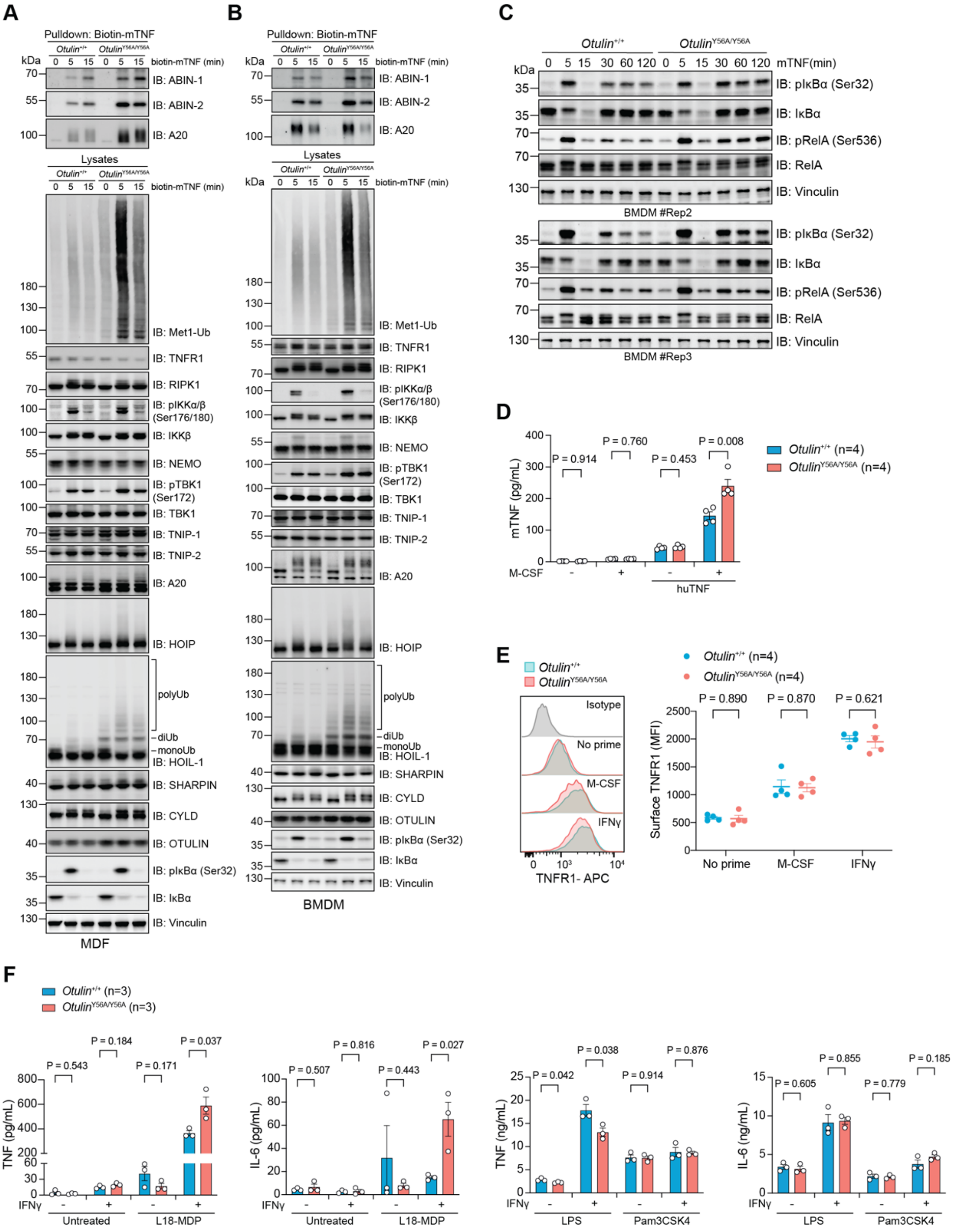


**Suppl. Figure S6. OTULIN-LUBAC interaction restricts TNF-induced NF-κB signalling.**

(A, B) Immunoblot analysis of TNF-RSC purified with biotin-mTNF and lysates from immortalised MDFs (A) and BMDMs (B) as described in Figures 3A, 3B.
(C) Immunoblot analysis of whole cell lysates from BMDMs stimulated with 10 ng/mL mTNF for the indicated time points.
(D) Cytokine concentrations in supernatants of BMDMs primed overnight with 20 ng/mL M-CSF and treated with 100 ng/mL huTNF for 24 hours.
(E) Flow cytometry analysis of surface TNFR1 levels in BMDMs of the indicated genotypes primed overnight with 20 ng/mL mIFN-γ or M-CSF.

(F) Cytokine concentrations in supernatants of BMDMs primed overnight with 20 ng/mL mIFNγ and treated with 200 ng/mL L18-MDP, 100 ng/mL LPS, or 100 ng/mL Pam3-CSK4 for 24 hours.

Representative results from three biological replicates (A, B) are shown. Data are presented as mean ± SEM and 'n' indicates the number of biological replicates in each group (D, E, F). Statistical analysis; multiple unpaired *t*-test (D), and two-way ANOVA with Turkey's multiple comparisons (E, F).

**Suppl. Figure S7**

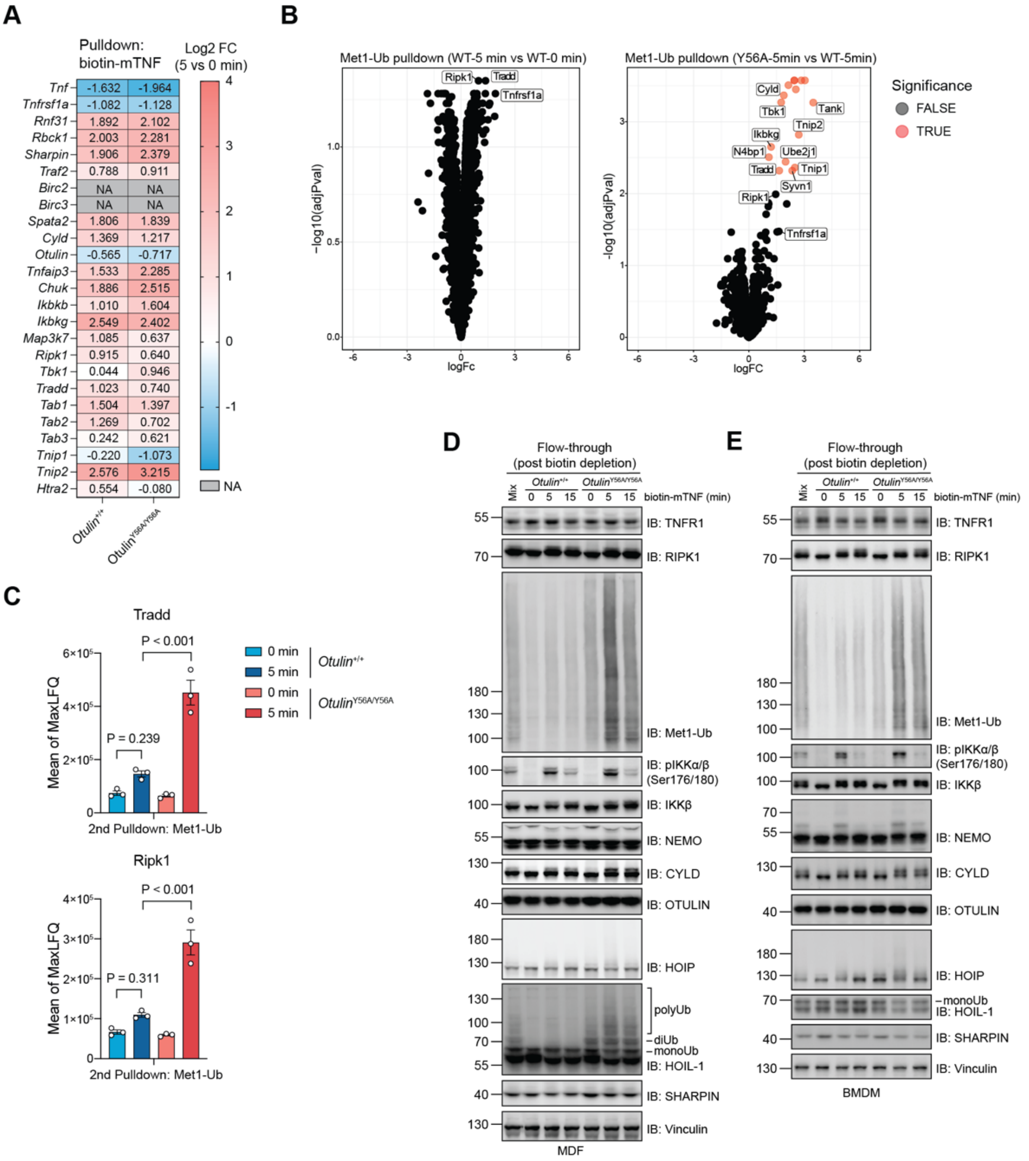


**Suppl. Figure S7. Disruption of the OTULIN-LUBAC interaction stabilises TNF-RSC after TNF dissociation.**

(A-C) Mass spectrometry analysis of TNF-RSC (A) and Met1-Ub-associated proteins in the flow-through fraction (B, C) from immortalised MDFs stimulated with 50 ng/mL biotin-mTNF shown as fold changes (A), volcano plots (B) and MaxLFQ intensities (C).

(D, E) Immunoblot analysis of HOIP-associated proteins in the flow-through fraction after depletion of biotin-mTNF. Data is related to Figures 4E, 4F.

Data are from three biological replicates (A-C) and one representative of two (D) or three (E) biological replicates. Data in (C) are presented as mean ± SEM from three biological replicates. Statistical analysis; two-way ANOVA with Turkey's multiple comparisons.

**Suppl. Figure S8**

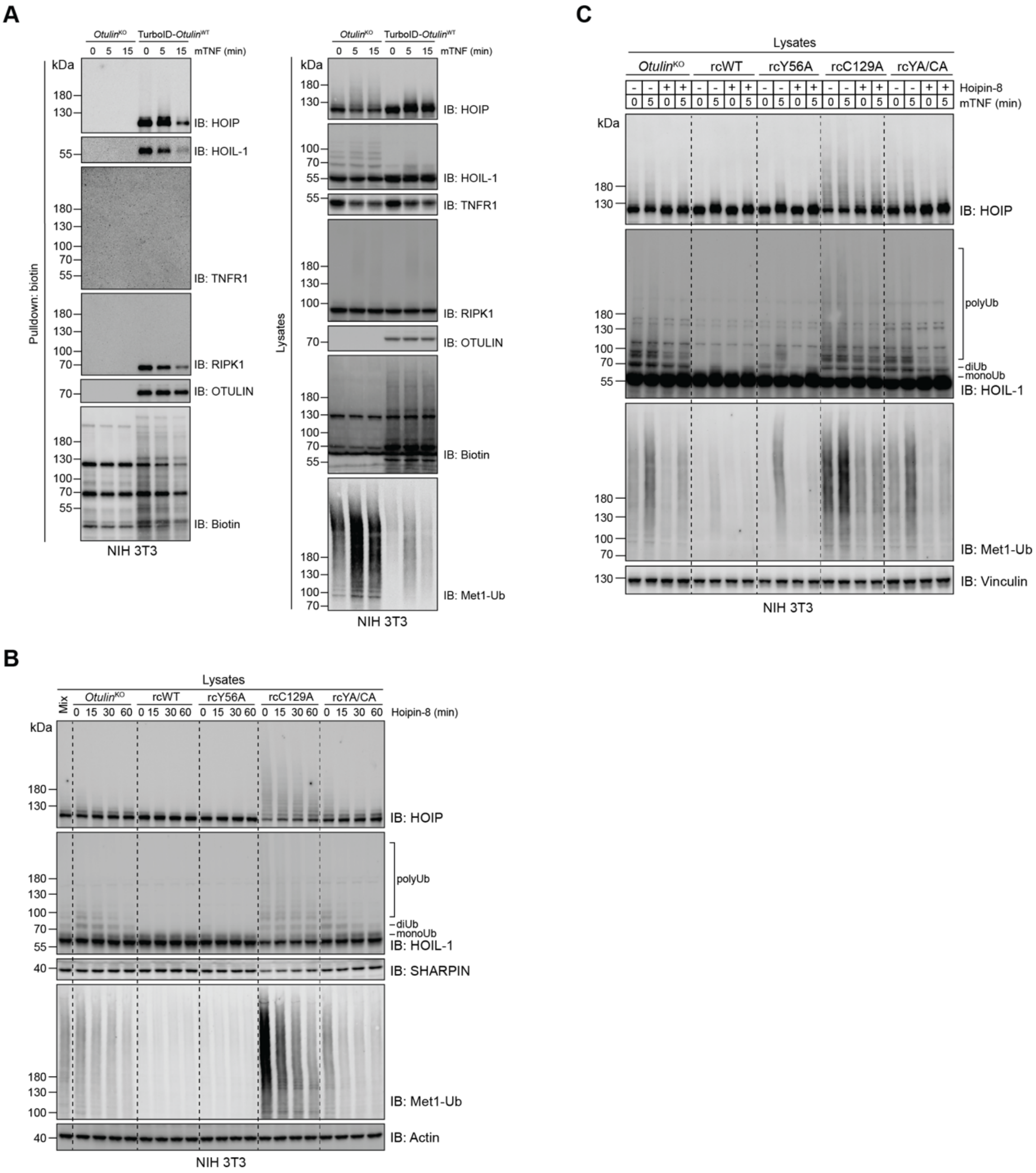


**Suppl. Figure S8. OTULIN counteracts LUBAC auto-ubiquitination by HOIP.**
(A) Immunoblot analysis of biotin pulldown fractions and lysates from *Otulin*$^{KO}$ or TurboID-OTULIN$^{WT}$ NIH 3T3 cells incubated with 100 µM biotin for 2 hours with or without 10 ng/mL mTNF stimulation at the last 5 or 15 minutes.
(B, C) Immunoblot analysis of lysates from *Otulin*$^{KO}$ NIH 3T3 cells reconstituted with OTULIN variants treated with 3 µM Hoipin-8 for the indicated time points (B) or for 30 minutes followed by stimulation with mTNF for 5 minutes (C).
Data are representative of two (A) and three (B, C) independent experiments with similar results.

**Suppl. Figure S9**

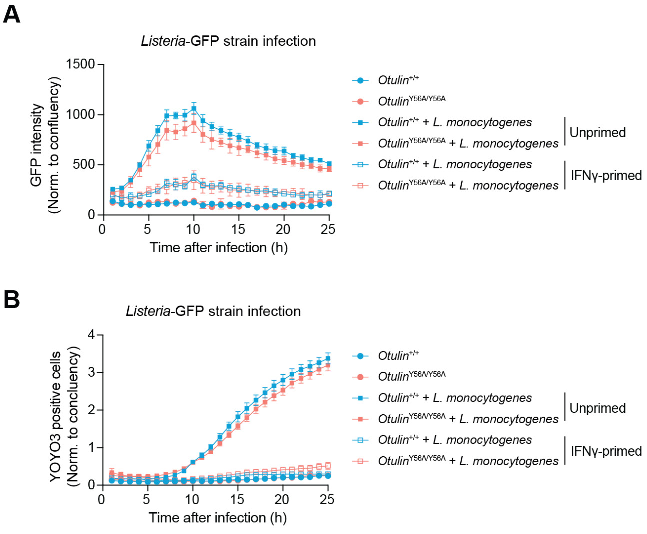


**Suppl. Figure S9. OTULIN$^{Y56A}$ mutation does not affect *Listeria* growth in BMDMs or cell death.**

(A) Intracellular proliferation of *Listeria*-GFP (MOI=1) in BMDMs primed overnight, or not, with 20 ng/mL mIFN-γ.

(B) Cell death of BMDMs in (a) measured by YOYO3 positivity and normalised to cell confluence.

Data are presented as mean ± SEM from three biological replicates.

### REFERENCES


1. Song, R. *et al.* A simple and rapid protocol for the isolation of murine bone marrow suitable for the differentiation of dendritic cells. *Methods and Protocols* **7**, 20 (2024).
2. Damgaard, R.B. *et al.* The deubiquitinase OTULIN is an essential negative regulator of inflammation and autoimmunity. *Cell* **166**, 1215-1230. e1220 (2016).
3. Kong, A.T., Leprevost, F.V., Avtonomov, D.M., Mellacheruvu, D. & Nesvizhskii, A.I. MSFragger: ultrafast and comprehensive peptide identification in mass spectrometry–based proteomics. *Nature Methods* **14**, 513-520 (2017).
4. Teo, G.C., Polasky, D.A., Yu, F. & Nesvizhskii, A.I. Fast Deisotoping Algorithm and Its Implementation in the MSFragger Search Engine. *Journal of Proteome Research* **20**, 498-505 (2021).
5. Yu, F. *et al.* Fast Quantitative Analysis of timsTOF PASEF Data with MSFragger and IonQuant. *Molecular & Cellular Proteomics* **19**, 1575-1585 (2020).
6. Yu, F., Haynes, S.E. & Nesvizhskii, A.I. IonQuant Enables Accurate and Sensitive Label-Free Quantification With FDR-Controlled Match-Between-Runs. *Mol Cell Proteomics* **20**, 100077 (2021).
7. da Veiga Leprevost, F. *et al.* Philosopher: a versatile toolkit for shotgun proteomics data analysis. *Nat Methods* **17**, 869-870 (2020).
8. Goeminne, L.J.E., Gevaert, K. & Clement, L. Experimental design and data-analysis in label-free quantitative LC/MS proteomics: A tutorial with MSqRob. *J Proteomics* **171**, 23-36 (2018).
9. Goeminne, L.J.E., Sticker, A., Martens, L., Gevaert, K. & Clement, L. MSqRob Takes the Missing Hurdle: Uniting Intensity- and Count-Based Proteomics. *Analytical Chemistry* **92**, 6278-6287 (2020).
